# “A novel Organ-Chip system emulates three-dimensional architecture of the human epithelia and allows fine control of mechanical forces acting on it.”

**DOI:** 10.1101/2020.08.02.233338

**Authors:** Antonio Varone, Justin Ke Nguyen, Lian Leng, Riccardo Barrile, Josiah Sliz, Carolina Lucchesi, Norman Wen, Achille Gravanis, Geraldine A. Hamilton, Katia C. Karalis, Christopher D. Hinojosa

## Abstract

Successful translation of *in vivo* experimental data to human patients is an unmet need and a bottleneck in the development of effective therapeutics. micro technology aims to address this need with significant advancements reported recently that enable modeling of organ level function. These microengineered chips enable researcher to recreate critical elements such as *in vivo* relevant tissue-tissue interface, air-liquid interface, and mechanical forces, such as mechanical stretch and fluidic shear stress, are crucial in emulating tissue level functions. Here, we present the development of a new, comprehensive 3D cell-culture system, where we combined our proprietary Organ-Chip technology with recent advantages in three-dimensional organotypic culture. Leveraging microfabrication techniques, we engineered a flexible chip that consists of a channel containing an organotypic epithelium surrounded by two vacuum channels that can be actuated to stretch the hydrogel throughout its thickness. Furthermore, the ceiling of this channel is a removable lid with a built-in microchannel that can be perfused with liquid or air and removed as needed for direct access to the tissue. The floor of this channel is a porous flexible membrane in contact with a microfluidic channel that provides diffusive mass transport to and from the channel. This additional microfluidic channel can be coated with endothelial cells to emulate a blood vessel and capture endothelial interactions. Our results show that the Open-Top Chip design successfully addresses common challenges associated with the Organs-on-Chips technology, including the capability to incorporate a tissue-specific extracellular matrix gel seeded with primary stromal cells, to reproduce the architectural complexity of tissues by micropatterning the gel, that can be extracted for H&E staining. We provide proof-of-concept data on the feasibility of the system using skin and alveolar epithelial primary cells and by simulating alveolar inflammation.

## INTRODUCTION

There is a rapidly growing interest in the development of 3D-organotypic cell based models, since these systems are more predictive of the human response to drugs and chemicals when compared to animal or 2D models^1,2,3^. Organs-on-Chips are microphysiological systems that support cell growth and differentiation and model important aspects of the spatial and chemical tissue microenvironment^4,5^. A central advancement in modeling the spatial organization of tissues came with the understanding of the role of biologically active components derived from the native extracellular matrix (ECM). It has been shown that functionalization and/or coating of artificial materials with ECM has profound effects on cellular survival, differentiation, and polarization^6,7^. Further advancements of *in vitro* cultures was achieved with the development of hydrogels made of naturally occurring ECM polymers, which form 3D scaffolds that enable organization of cell structures reminiscent of the native tissues and support organ-level functions^8,9,10,11^. Despite the great potential of 3D hydrogel culture models^11,12^, especially for multicellular human primary biological systems such as organoids^13^, 3D cultures have several important limitations including complex sample handling, high cost, and challenges with penetration of compounds and access to the lumen in organs such as the intestine for barrier function or transporter studies^14,15^. These systems also lacked an important component that is a critical driver of differentiated function *in vivo*, dynamic microenvironment that includes flow, mechanical forces, and interactions with both resident and circulating immune cells. The limitations associated with the 3D hydrogel cultures led to the development of the engineered microphysiological systems^16^. Along these lines, several groups have adapted microfabrication methods derived from the computer microchip industry to produce micrometer-scale systems that can be used to culture human cells^17^. These microengineered systems, referred to as Organs-on-Chips, consist of two or more microfluidic channels, separated by a semi-permeable porous material able to accommodate different types of cells in structures resembling the tissues of origin^18,19^. In addition, with the use of specific ECM coatings, flow regimes, and other special conditions such as stretching and hypoxia, they are currently the most advanced platforms available for drug safety and efficacy studies^20,21,22^. However, most Organs-on-Chips designs do not include stroma, a functional component of the epithelial and parenchymal organs containing different types of mesenchymal cells^23,24^ and one that becomes critically important as move towards disease modeling, more complex tissues, and more complex biological mechanisms. Incorporation of a relatively large gel volume enables inclusion of enough mesenchymal (fibroblast, smooth muscle) cells to obtain epithelial to mesenchymal ratios as required for recapitulation of the associated biological effects *in vitro*. As natural ECM hydrogels have stiffness closer to the organs *in vivo*, they provide better support for a physiologically relevant cell behavior and gene expression^25^. Here we describe the development of a newly designed chip that addresses this need by enabling the inclusion of a relatively large volume of hydrogel with embedded mesenchymal cells to recapitulate the stromal component, overlaid with epithelial cells and supported by a vascular component. We also provide proof of concept of how we leveraged this new chip, referred to as the Open-Top Chip, to culture two different epithelial cell types – skin and alveolus– which represent examples of stratified and simple epithelium, respectively. Finally, we present the response of the alveolar epithelium to a common experimental infectious challenge, to highlight a potential application.

## METHODS & MATERIALS

### Fabrication and assembling of the Open-Top Chip

All the constituent parts of the Open-Top Chip, as well as the tooling to mold the individual components, were designed using SolidWorks. The platform comprises of four parts made of polydimethylsiloxane (PDMS): a bottom spiraled shaped microfluidic channel which has a cross-sectional profile of 400μm (height) × 600μm (width), a porous membrane which has a thickness of 50μm with pores 7μm diameter, spaced in a 40μm hexagonal packed geometry, a channel which has a diameter of 6mm and an height of 4mm, and a channel lid that includes a microfluidic channel. The channel is round to equilibrate the tension forces acting on the hydrogel-wall interface and to eliminate geometric stress concentrators that would occur at the vertices of a rectangular shaped wall, which in prototype designs resulted in delamination of the hydrogel from the PDMS wall. The bottom spiraled shaped microfluidic channel, channel and lid was fabricated using soft lithography/replica molding of polydimethylsiloxane (PDMS) from a mold created with high resolution SLA 3D printing (Protolabs). The membrane was fabricated using previously published methods^26^. Briefly, membranes are made by casting PDMS over a silane coated silicon wafer master, which contains a post-array formed using photolithography techniques. The weight ratio of PDMS base to curing agent was 10:1. During the curing of the PDMS at 65°C overnight, constant pressure was applied using a custom built pneumatic press that applies a pressure of 40PSI (276KPa) to the master to ensure intimate contact and penetration of PDMS throughout the silane coated master. This process produced 50μm thick PDMS membranes with circular through-holes. The PDMS parts, bottom spiraled shaped microfluidic channel, membrane and channel were manually aligned and, then, bonded together by oxygen plasma treatment (30W for 30 seconds at 335mtorr using a Femto Science Covance Oxygen plasma machine). Irreversible bonding was achieved by curing the PDMS parts at 60°C overnight.

### Pneumatic actuator

Pneumatic actuation of the Open-Top Chip vacuum channels was performed by using a custom programmable vacuum pressure regulation system. The system consisted of an electro-pneumatic vacuum regulator (ITV0091-2BL, SMC Corporation of America) controlled electronically by an Arduino Leonardo and MAX517 digital to analogue converter. The electro-pneumatic vacuum regulator generates a sinusoidal vacuum profile with amplitude and frequency tunable by the user. Cyclic strain ranging from 0 to 15% was generated by applying negative pressure to the vacuum channel of the Open-Top Chip at an amplitude ranging from 0 to −90kPa and frequency of 0.2 Hz.

### Peristaltic pump

Perfusion of the Open-Top Chip was achieved with a multichannel programmable peristaltic pump (Ismatec IPC ISM934C; Cole-Palmer: EW-78001-42) using Click’n’go™ cartridges (Ismatec Click-N-Go Cassette/Cartridge, 2-Stop, POM-C; Cole-Palmer: EW-78001-95) for two-stop tubes (Ismatec Pump Tubing 2-Stop, PharMed^®^ BPT, 0.25 mm ID; Cole-Palmer: EW-95723-12). Two-stop tubes were connected to the Open-Top Chip inlet ports by 19-gauge hypodermic tubing (1inch) (MicroGroup: 316H19RW) and linked to the medium reservoirs through an 18-gauge hypodermic tube. The fluidic system comprises of two inlet reservoirs consisting of 15 ml conical tubes capped with a lid fabricated to allow the insertion of 18-gauge hypodermic needles (4inch) (MicroGroup: 316H18RW) and linked to the top and bottom Open-Top Chip fluidics. Two separate Masterflex Transfer Tubing, PharMed^®^ BPT, 1/32” ID × 5/32” OD (Cole-Palmer: EW-96880-02) lines connected to each chip inlet ports by 18-gauge hypodermic tubing (1inch) (MicroGroup: 316H18RW), Open-Top Chips, peristaltic pump and two outlet reservoirs consisting of uncapped 15 conical tubes held by a rack. Media were routed from the inlet medium reservoirs through the Open-Top Chip across the peristaltic pump into the outlet reservoirs. The pump, reservoirs, and chips were places in a standard tissue culture incubator that provided the proper carbon dioxide concentration to buffer the pH of sodium bicarbonate buffered media contained in reservoirs. The peristaltic pump setup enables the parallel perfusion of 12 Open-Top Chips, with most incubators fitting two setups to enable the culture of 24 chips per incubator.

### Chemical activation

Chemical activation of the PDMS channel is an important step to ensure the covalent cross-linking of the ECM polymer to the PDMS and to guarantee the adhesion of the hydrogel throughout the entire duration of the culture. Briefly, ER-1 (Emulate reagent: 10461) and ER-2 (Emulate reagent: 10462) are mixed at a concentration of 1mg/ml and added to the bottom microfluidic channel and to the channel. The platform is then irradiated with high power UV light having peak wavelength of 365nm and intensity of 100μJ/cm^2^ for 20min using a UV oven (CL-1000 Ultraviolet Crosslinker AnalytiK-Jena: 95-0228-01). The procedure is repeated twice to maximize the activation of the surface.

### Stamp for micropatterning gels

The micropatterning stamps were designed using SolidWorks software and 3D printed in a non-cytotoxic resin (MicroFine Green or Somos^®^ WaterShed XC 11122 resin, Proto Labs) with 15μm layer resolution. The cylindrical 3D printed stamps were used to micromold the hydrogel surface to specific heights and geometries (2mm for skin and 200μm for alveolus) leveraging the stamp design. We opted to use a flat stamp design to reduce the irregularity of the gel surface which can lead to optical distortions and complicate the imaging of the epithelium channel. The micropatterning technique involved four basic steps: first, remove the top lid to deposit the hydrogel into the channel by pipetting; second, introduce the micropatterned stamp over the hydrogel into the channel; third, patterning the gel by allowing the hydrogel polymerization with the stamp inserted into the channel; and forth, remove the stamp and aspirate the excess of hydrogel. The surface of the stamps is undersized 100μm to permit the escape of air during the casting operation, which can otherwise trap bubbles into the hydrogel. We take advantage of the openable Open-Top lid to introduce these specially designed micropatterned cylindrical stamps which imitate the topography of the natural human epithelium-stroma interface to cast hydrogel surface at micrometer scale or to form a flat gel surface which are often convenient over complex geometries because flatness facilitates the spatial confinement of the epithelial layer and makes imaging cells easier.

### Cell expansion

Human Primary Alveolar Epithelial Cells (Cell Biologics: H-6053), Lung Fibroblasts (Cell Biologics: H-6013), Normal Human Lung Smooth Muscle Cells (Lifeline:FC-0046), Normal Human Lung Fibroblasts, Primary (Lifeline: FC-0049) and Human Lung Microvascular Endothelial Cells (Lonza: CC-2527) were grown in T-75 culture flasks in an atmosphere of 5% CO_2_ at 37°C according to the instructions provide by the manufacturers. Briefly, Primary Alveolar Epithelial Cells (P1) were cultured in SAGM medium (SAGM Lonza: CC-4124). Lung Fibroblasts (P1), Normal Human Lung Smooth Muscle Cells (Lifeline:FC-0046), Normal Human Lung Fibroblasts, Primary (P1) were cultured in DMEM/F-12 (GIBCO: 11320082) containing 10% Heat Inactivated HyClone™ FetalClone™ II Serum (U.S.) (GE Healthcare Life Sciences: SH30066.03) and 1% penicillin-streptomycin (GIBCO: 15140122) and 1% GlutaMAX (GIBCO: 35050061). Human Lung Microvascular Endothelial Cells were cultured in EGM-2MV (Lonza: CC-3202). Primary Epidermal Keratinocytes; Normal, Human, Neonatal Foreskin (HEKn) (ATCC^®^: PCS-200-010™), Primary Dermal Fibroblast Normal (HDFn) (ATCC^®^: PCS-201-010™) and Human Microvascular Endothelial Cell (ATCC^®^: CRL-3243™) were grown in T-75 culture flasks in an atmosphere of 5% CO_2_ at 37°C according to the instructions provide by the manufacturers. Normal Human Neonatal Foreskin (P1) were cultured in Keratinocyte-SFM (1X) medium (Thermo: 17005042). Primary Dermal Fibroblast Normal (P1) were cultured in DMEM/F-12, GlutaMAX™ supplement (GIBCO : 10565018) containing 10% Heat Inactivated HyClone™ FetalClone™ II Serum (U.S.) (GE Healthcare Life Sciences: SH30066.03) and 1% penicillin-streptomycin (GIBCO: 15140122) and 1% GlutaMAX (GIBCO: 35050061). Human Microvascular Endothelial Cells were cultured in EGM-2MV (Lonza: CC-3202).

### Stromal compartment

Following the chemical activation of the Open-Top Chip and chip washing with ER2 (2×200μl), a stroma equivalent ECM mix was prepared using a solution of bovine type I collagen (Advanced BioMatrix 5133). Briefly, 8 volumes of collagen I gel solution (10mg/ml) were mixed with 1 volume of 10X EMEM (Lonza: 12-684F) and 1 volume of 10X Reconstruction buffer(Anacker and Moody 2012) while kept on ice to obtain a collagen solution at a final concentration of 8.0 mg/ml. Typically, 10μl of 1 N NaOH for ml of collagen mixture were added to adjust its pH at around 7.4. Following the addition of NaOH the collagen mixture was thoroughly mixed. The change to the proper pH of the collagen mixture was visually evaluated by the switch in color from yellow to a brown/light pink. At this point, 1 volume of fibroblast (and/or Smooth muscle cells) solution (0.5 × 10^6^ cells/ml) was added to the neutralized collagen solution and fully mixed by pipetting up and down without introducing bubbles.

The collagen I hydrogel with embedded fibroblasts and/or smooth muscle cells was pipetted in the stromal compartment of the Open-Top Chip, compressed using a flat stamp to 2mm (skin) or 200μm (alveolus) tall gels and let polymerize at 37⍰°C, 5% CO_2_ for 60⍰min. As per the manufacturer, the composition of the described hydrogel gives an average stiffness of around 950 Pa (and Young Modulus (E) of around 4KPa) that partially matches both that of the lung (stiffness 0.9 – 2.5kPa^27^ and elastic modulus (E) 0.44 to 7.5kPa^28^) and that of the normal skin (Young Modulus (E) 0.1 −10KPa^29^).

After polymerization, the stamp was removed and the hydrogel surface and bottom spiraled channel were coated with a solution of collagen IV for skin and collagen IV, fibronectin and laminin (1200μl of 1mg/ml solution + 300μl of 1mg/ml solution + 30μl of 1mg/ml) for alveolus. These coatings simulate the basal lamina of the respective epithelia and improve epithelial cell adhesion. ECM coating solutions were left to polymerize for 2-4 hours at 37⍰°C, 5% CO_2_ to ensure that the spiraled channel and the epithelial surface of the stroma equivalents were properly coated. After coating, the recreated stroma was maintained in a static condition. Both top and bottom channels were filled with medium, DMEM/F12 GlutaMax supplemented with 10% serum (Heat Inactivated HyClone™ FetalClone™ II Serum (U.S.) GE Healthcare Life Sciences Cat #: SH30066.03) and kept in this submerged status for 24/48 hours to let fibroblasts and smooth muscle cells recover from the embedding procedure.

### Epithelial channel

After Collagen I hydrogel gelation and ECM coating, the recreated stroma is typically seeded with epithelial cells on the top and cultured in a submerged state until the cells form a compact monolayer. Epithelial cells are seeded side and allowed to sediment directly against the ECM gel. Upon attachment of the epithelial cells, the stroma (which is crosslinked to the PDMS) and epithelium form a tight barrier which balance the hydrostatic pressure exerted by the medium in the bottom. The monolayer lining the top of the stomal equivalent, together with the top microfluidic channel, forms what we define the epithelium channel. The top microfluidic channel is connected to an inlet and an outlet port that can be used to perfuse the epithelial tissue with culturing medium, airflow, or to expose it to static air. This feature is particularly important because it enables the establishment of an air-liquid interface (ALI). The top microfluidic channel can also be opened or removed to allow direct access to the stromal compartment (open-top capability). Once the epithelial barrier is established, the bottom channel can be perfused without the risk of medium leaking into the epithelium channel, and it is possible to establish the air-liquid interface that is essential for many epithelia to fully differentiate.

### Open-Top Chip microfluidics and vascular channel

Following the establishment of the epithelial barrier, endothelial cells are seeded inside the bottom channel and left to colonize its entire internal wall surface to form a confluent tube whose lumen interconnects with the inlet and the outlet making it accessible to perfuse medium, blood or other fluids (serum, medium with PBMC, blood surrogate) as per the experimental needs. The application of flow supported establishment of the barrier and induced maturation of the tight junctions. When the bottom microfluidic channel is lined with endothelial cells during the culturing phase, it forms what we define as the vascular channel. The spiraled section of the bottom channel interacts directly with the basal side of the recreated stroma by the PDMS semi-permeable porous membrane. When this membrane is populated with endothelial cells, it emulates the stoma-capillary interface where exchange of nutrients and waste products between organ cells and the circulatory system occurs. The PDMS porous membrane has the structural function to physically contain the hydrogel in place and to feed and hydrate the recreated stroma and the epithelium with medium. We opted for a spiraled shaped channel to maximize the area of fluid (medium) in direct contact with the circular section of the stromal compartment, and to simultaneously maintain constant laminar flow along the entire length of the channel. As a result, the wall-shear rate along the entire length of the endothelial channel can be modified by simply changing the flow rate (Fig. S1). Primary human endothelial cells are cultured on all surfaces of the bottom microfluidic channel. The full channel coverage is achieved by chemically activating the PDMS surface of the microfluidic channel wall and by coating it with ECM to improve endothelial cell adhesion and coverage. Following the ECM coating a double seeding technique is used to form a complete endothelial lumen. The double seeding method is a two-part process, the first part involves the loading of the chip bottom microfluidic channel with an endothelial cell suspension of 3 × 10^6^ cells/ml. The chip is then flipped upside down to let endothelial cells attach to the membrane side (top/ceiling wall) of the microfluidic channel during a one-hour incubation. The second part involves flushing the bottom microfluidic channel with medium to remove cellular debris and a second loading of the microfluidic channel with an endothelial cell suspension of 3 × 10^6^ cells/ml. During this second seeding the chip is not flipped and set flat to let endothelial cells attach to the bottom wall of the microfluidic channel during a one-hour incubation, which is once again flushed to remove cellular debris. After flushing, the chip is kept into the incubator in static condition for a few hours to overnight to let endothelial cells proliferate and form a complete lumen before connecting it to the pump.

### LPS-induced inflammation

Experiments to confirm the functionality of the vasculature channel and its ability to respond to stimuli were conducted by exposing the epithelial channel of the Open-Top Organ-Chip to 50μl of 10μg/ml Lipopolysaccharide (LPS)(Escherichia coli O111:B4, Sigma L4391) solution for 8 hours. After the treatment, the vascular channel was either stained for ICAM-1 or perfused with CD41 labelled platelet whole blood.

### Immunostaining

Open-Top Organ-Chips were fixed in 10% neutral buffered formalin. Immunofluorescence staining of the endothelial and epithelial cells was performed by flowing the reagents through their respective microfluidic channel. Alternatively, the epithelium equivalents (recreated stroma with epithelium) were extracted from the chip, fixed in 10% neutral buffered formalin for at least 24 hours, dehydrated by a series of alcohol washes of increasing concentrations (70%, 80%, two of 95%, three of 100% ethanol), cleared by two washes in xylene and infiltrated by paraffin using a tissue processor (Leica TP1020) before being embedded into paraffin wax (Leica HistoCore Arcadia H & C) and cut in sections (typically between 8-12 micrometers thick) with a microtome (Leica: RM2235). Sections were placed on Poly-L-Lysine coated microscope glass slides (Sigma: P0425) according to standard immunohistochemistry procedures and processed for immunohistochemistry analysis. For H&E staining, paraffin was removed by two changes in xylene of 5 minutes, the slides were rehydrated by sequential ethanol washes (two of 100% ethanol, and one wash each of 95% and 70% ethanol) and a final rinse in ddH_2_O for 2 minutes. Slices were then stained in Gill 2 hematoxylin solution (VWR: 10143-146) for 10 min, washed sequentially with deionized water for one minute, washed with acetic alcohol for 20 seconds, dipped in bluing agent (VWR: CA95057-852) 10 times, rinsed with 70% ethanol for one minute, before being counterstained with eosin Y alcoholic (VWR: 10143-132) for 2 minutes. This was followed by a series of ethanol washes (two of 95% ethanol, three of 100% ethanol), and a final clearing by two washes in xylene for 5 minutes. Slides were then mounted with Permount (EMS: 17986-05) and glass coverslips. For formalin-fixed paraffin-embedded tissues, antigen-retrieval was performed by boiling the tissue in epitope retrival solution (IHC-Tek: IW-1100) using a pressure cooker (IHC-Tek: 1102). For immunofluorescence staining cells were permeabilized using 0.1% Triton X-100 (Sigma: T8787) solution in Dulbecco’s phosphate-buffered saline for 20 min, followed by blocking of non-specific binding with blocking solution containing 1% bovine serum albumin (Thermo: 37525), and 5% normal donkey serum (Millipore: S30-100ML) in Dulbecco’s phosphate-buffered saline. After blocking, epithelium equivalents were incubated with primary antibodies (1:100) at 4⍰°C overnight, followed by washing steps and sandwich labeling with appropriate secondary antibodies (1:200) at 4⍰°C overnight, counterstained with DAPI (Thermo: D1306) and imaged. The primary antibodies used included MUC5AC (Thermo: PA5-34612), beta IV Tubulin (Abcam: ab11315), Isoforms TA p63-α, −β, −γ (Biolengend: 618902), CC16, Human, mAb AY1E6 (HyCultBiotech: HM2178), ABCA3(Abcam: ab24751), LAMP3 (Abcam: ab111090), Podoplanin (AT-1α) (Abcam: ab128994), surfactant B (Abcam: ab40876), HTI (Terrace Biotech: TB-29AHT1-56), HTII (Terrace Biotech: TB-27AHT2-280), E-Cadherin (Abcam: ab1416 and ab40772), ZO-1 (Thermo: ZO1-1A12) conjugated Alexa Fluor 647 (Invitrogen: 33-9100-A647), Cytokeratin 14 (Abcam: ab7800), Cytokeratin 10 (DE-K10) (Thermo: MA5-13705), Involucrin (SY5) (Thermo: MA5-11803), Filaggrin (FLG01) (Thermo: PA5-79267), VE-cadherin (Abcam: ab33168), PCNA (Thermo: PA5-32541), PECAM-1 (Abcam: ab9498), VWF (Abcam: ab8822) and surfactant C (Seven Hill: WRAB-9337). Donkey Anti-Mouse IgG H&L (Alexa Fluor^®^ 488) (Abcam: ab150105), Donkey Anti-Mouse IgG H&L (Alexa Fluor^®^ 568) (Abcam: ab175472), Donkey Anti-Mouse IgG H&L (Alexa Fluor^®^ 647) (Abcam: ab150107), Donkey Anti-Rabbit IgG H&L (Alexa Fluor^®^ 488) (Abcam: ab150073), Donkey Anti-Rabbit IgG H&L (Alexa Fluor^®^ 568) (Abcam: ab175470), Donkey Anti-Rabbit IgG H&L (Alexa Fluor^®^ 647) (Abcam: ab150075) and Goat anti-Mouse IgG1 Alexa Fluor 568 (Thermo: A-21124) and anti-Mouse IgM Alexa Fluor 488 (Thermo: A-21042) were used as secondary antibodies. Blood was labelled with CD41 Monoclonal Antibody (VIPL3), PE (Thermo: MHCD4104) by direct addition of the antibody (1:100) into the blood and incubated for 15min at RT.

### ELISA immunoassay

PBS (200μl) was washed through the epithelial surface of the Open-Top Alveolus-Chip by pipetting, and collected, for measurement of Surfactant C levels using an ELISA kit for Human Surfactant Protein C (LifeSpanBioScience: LS-F12438), following the manufacturer’s protocol. The color reaction was measured at 450nm with a Synergy NEO HTS Multi-Mode microplate reader (BioTek). Samples were stored at −80°C until analysis. Similarly, the presence of cytokines (IL-6, IL-8 and MPC-1) was detected from medium samples by Human TLR-induced Cytokines II: Microbial-induced Multi-Analyte ELISArray Kit (Qiogen: MEH-008A) following the manufacturer’s protocol. The color reaction was measured at 450nm with a Synergy NEO HTS Multi-Mode microplate reader (BioTek). Samples were stored at −80°C until analysis.

### Microscopy and image analysis

All bright field videos were acquired using a Zeiss Observer.Z1 inverted microscope equipped with ORCA-Flash 4.0 digital camera (Hamamatsu: C11440), EC Plan Neoflaur 10X/0.3NA objective, 20X/04NA objective, 40X/0.6NA Corr Ph2 M27 objective, and ZEN software. Immunostaining were captured with fluorescence microscope Olympus U-LH100-3 microscope equipped with ORCA-Flash 4.0 digital camera (Hamamatsu, C11440) using a LUC PL FLN 40X/0.6, 20X/04NA or 10X/0.3NA objective and CellSense software. Confocal image and video co-localization, and quantitative analysis (intensity measurements) were performed using ImageJ software.

### Statistical analysis

To assess statistical significance of differences between groups, we used unpaired t-Test with Welch’s correction, Two-way ANOVA, and the Turkey honestly significant different (HSD) test using correlation analysis in the GraphPad Prism software program (San Diego, CA). The significance of the correlations between measurements was determined based on the null hypothesis that R=0. Significance was defined as α<0.05. Measurement are reported as means ± SEM and significant p values: *<0.05, **<0.01,***<0.001,****<0.0001.

## RESULTS

### Open-Top Chip design: technical features

We engineered the Open-Top Chip assembly to have a 35⍰mm⍰×⍰17⍰mm footprint and be comprised of four parts: the bottom spiraled shaped microfluidic channel, the membrane, the circular stromal and epithelial channel, and the lid that can serve as a microfluidic channel as well (Fig. 1). The channel has a tissue culture area of 0.32cm^2^ hosting the hydrogel that is chemically crosslinked to the PDMS wall and membrane, before being overlaid with epithelial cells. Chemical activation of the PDMS channel is an important step to ensure robust hydrogel-polymer adhesion for long-term cell cultures. Once the circular channel was chemically activated, the hydrogel, containing embedded primary fibroblasts and/or smooth muscle cells, can be introduced in the Open-Top Chip to cross-link with the channel and form the stroma equivalent. The surface area of the bottom spiraled microfluidic channel directly interfaces with the tissue culture area of the channel is 0.18cm^2^. The porous flexible PDMS membrane (2.5% porosity), interposed between the spiraled microfluidic channel and the circular channel, provides a supporting structure for the organotypic culture, while allowing diffusion transport of nutrients, metabolites or experimental compounds (Fig. 1).

**Figure 1.**
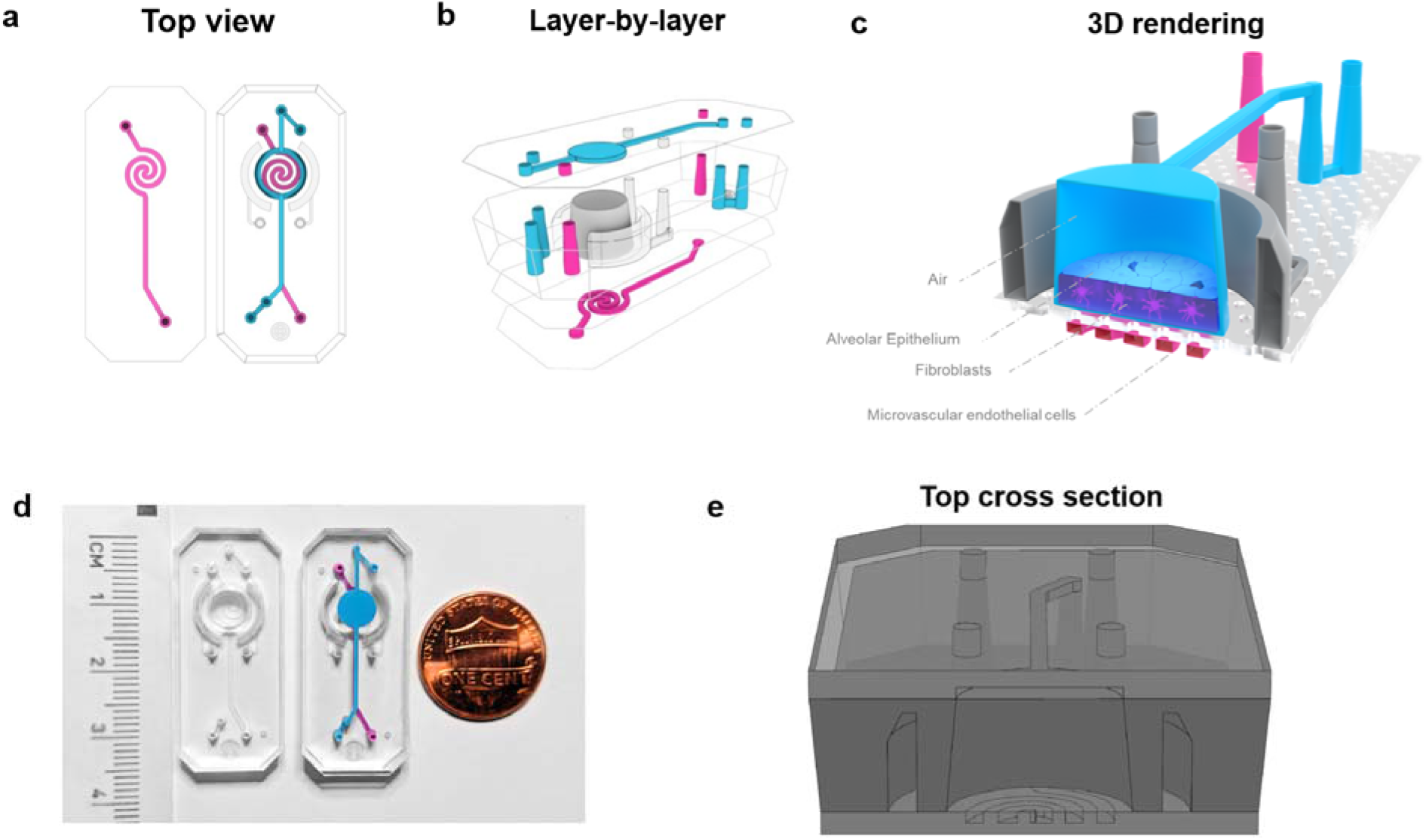
Schematic of the Open-Top Chip design: (a) Top-view projection and (b) layer-by-layer view of the Open-Top Chip showing the relative position of the upper fluidic channel (cyan), the two curved vacuum-channels positioned alongside the stretchable stromal compartment (grey) and the lower spiraled channel (magenta). (c) Tridimensional rendering illustrating the Open-Top Alveolus-Chip and the spatial orientation of three biological elements: the alveolar epithelium lining the upper surface of the recreated stroma covered by upper fluidic channel; the tridimensional recreated stroma located underneath the alveolar epithelium and encapsulated between the channel wall and the porous-elastic membrane; the endothelium lining the lower spiral-fluidic channel and the porous-elastic membrane forming the vascular channel. (d) Open-Top Chip filled with colored fluid to illustrate the relative position of the top (cyan) and bottom channel (magenta); size comparison illustrating the dimensions of the Open-Top Chip compared to a familiar object (penny). (e) Top cross section illustrating Open-Top Chip channel dimensions and overall design.

### Pneumatic stretching

The two semi-circular shaped hollow vacuum channels surrounding the stromal compartment can be subjected to negative vacuum pressure that deforms their walls, and subsequently the hydrogel attached to those walls. This process is referred to as pneumatic stretching, or simply stretching. The stretching range was chosen based on a quasi-linear dependence between the amounts of the vacuum pressure and magnitude of strain applied. The custom programmable vacuum pressure regulation system can modify the stretch frequency in the range of 0.01 to 0.5Hz, which generates stretching/relaxation cycles ranging from 2 to 100 seconds. When the negative vacuum pressure is decreased, the PDMS returns back (elastic recoil) to its original shape and relaxes the strain exerted on the stromal compartment as shown (Movie 2.1; Movie 2.2). In living organs, both the stroma and epithelium experience mechanical stretching and with this design it becomes possible to model motion such as breathing or peristalsis. Application of negative pressures in the range of 0 to −90KPa by our vacuum pressure regulation system generated a dynamic range of strain in the range 0 to 15% (Fig. 2 a), with the maximum and minimum strains detected respectively at the external edge and at the center of the channel (Fig. 2 b). To estimate the strain applied, fluorescent beads were embedded in the gel and the distance between two fluorescent beads at given heights within the gel was measured optically in the presence or absence of vacuum. Thus, the strain is calculated by the difference between the two distances as measured with and without vacuum, divided by the distance when no vacuum was applied.

**Figure 2.**
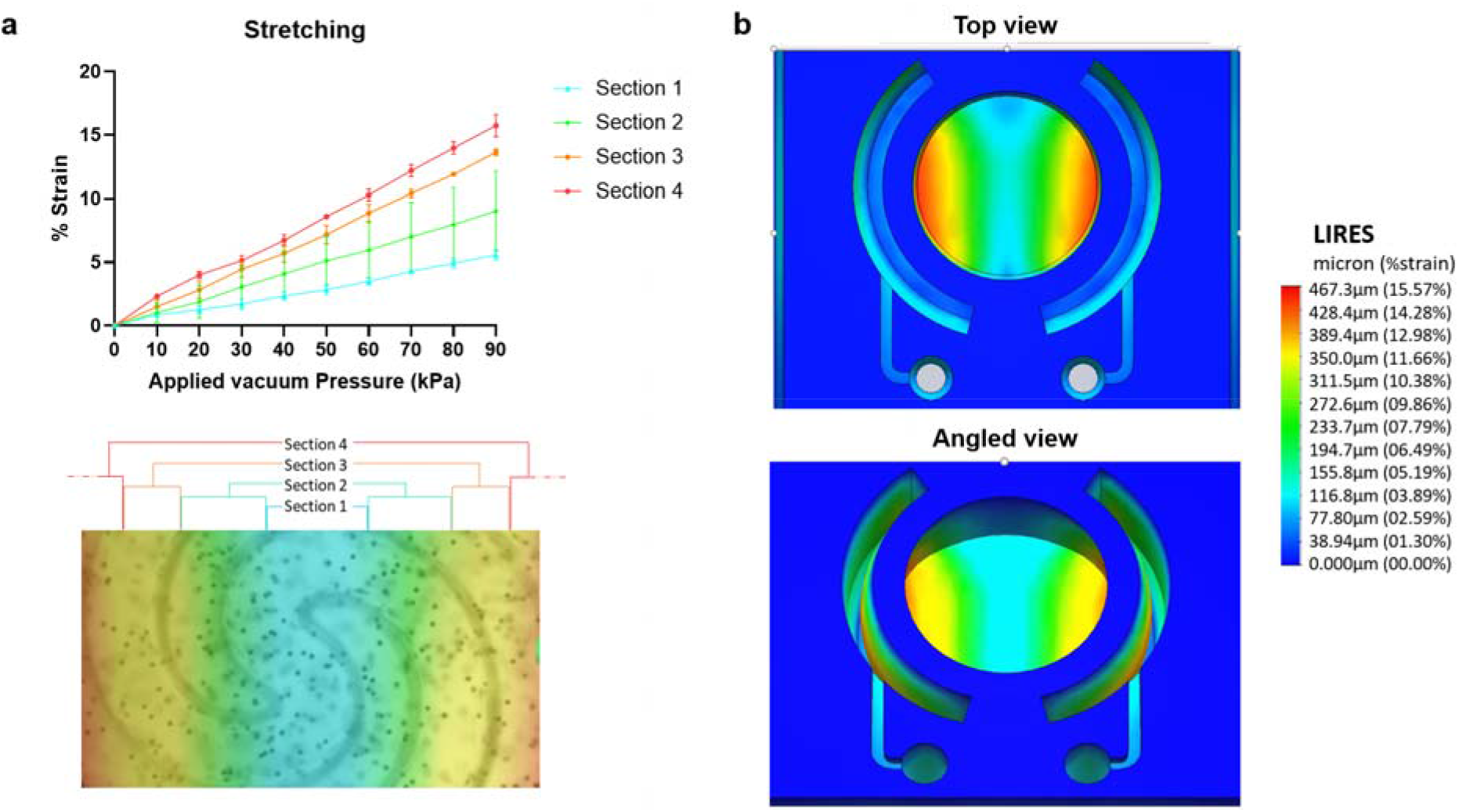
Open-Top Chip stretching: (a) Graph showing the correlation between negative pressure applied to the two curved vacuum-channels and percent of strain (top image); color code map illustrating the area of relative stretching measured as latex beads displacement respect to initial length (% strain) on the surface of the stroma equivalent (bottom image). (b) Finite Element Analysis (FEA) modelling the applied mechanical stretching along the surface of the stroma equivalent (top and angled view).

### Stamp for micropatterning

Micropatterning of extracellular matrix (ECM) hydrogel is used to replicate the spatial configuration of tissue interface microarchitecture, enabling study of the cellular response to irregular ECM geometries. The process of micropatterning by stamping was inspired by lithography, a process used to transfer geometric patterns from a wafer stamp to a soft substrate. Similarly, we can engineer the surface of a wafer stamp made of biocompatible materials to transfer a cell-size geometric pattern to an ECM substrate, as described in detail in methods. The diameter of the stamp is undersized in respect to the channel diameter to permit escape of air during the casting operation and thus prevent trapping of bubbles into the hydrogel. The stamp can be patterned with positive or negative micro-scale texture, such as pillars or holes with different geometrical packing or free style/random pattern (Fig. 3).

**Figure 3.**
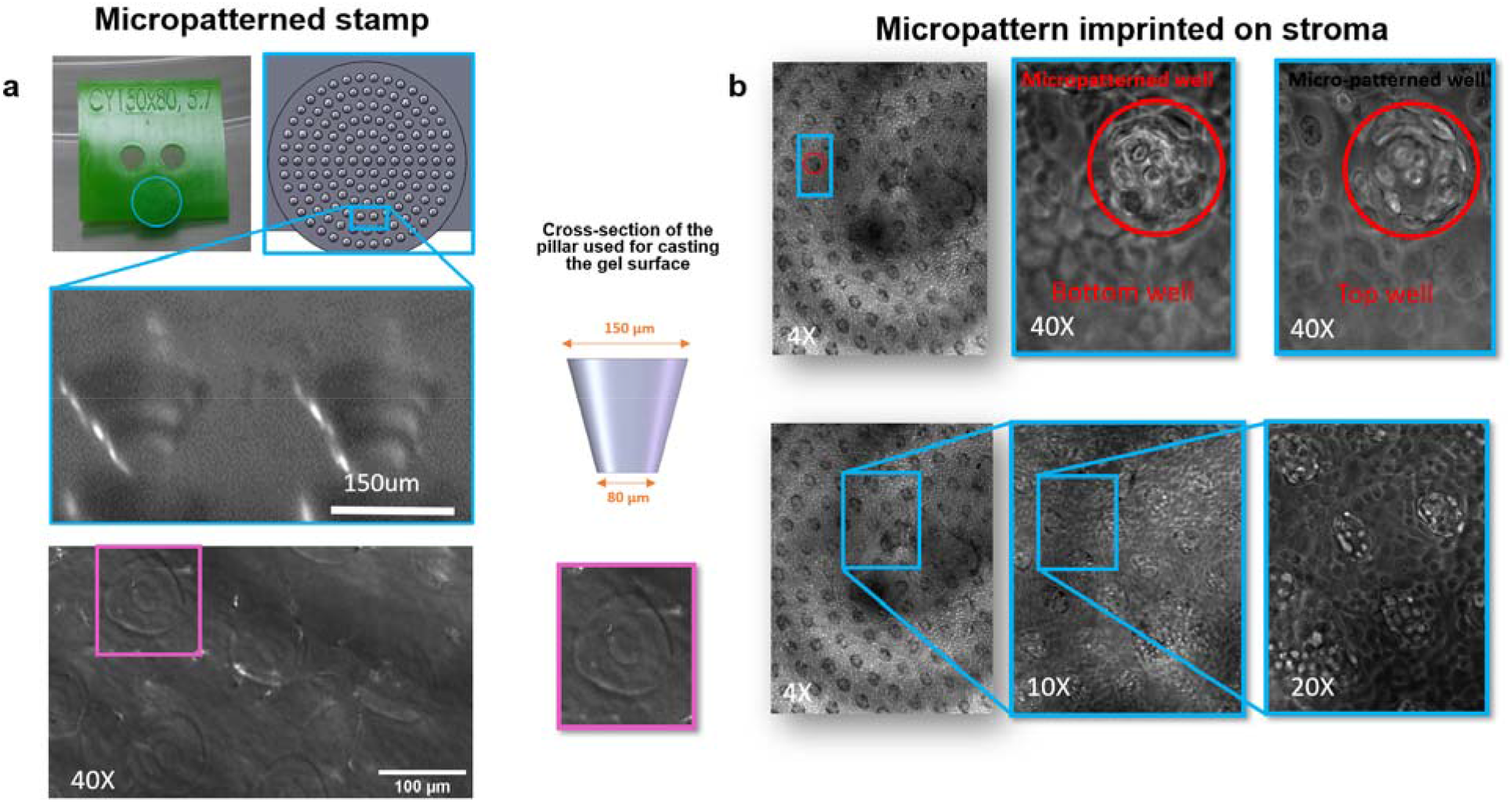
(a) Stamp micropatterned with a conical 150μm×80μm μm pillar design 3D printed in MicroFine Green by Protolab (top image). Image illustrating pillar design after 3D printing (middle image) and phase contrast image showing the marks that the stamp (150μm×80μm pillar design) leaves on the recreated stroma. (b) Phase contrast images of the stroma-epithelial interface illustrating a micropatterned stroma lined with epithelial cells at different magnifications; The red circles show a micropatterned well at two different heights.

### Endothelial cell incorporation to build the vasculature channel in the Open-Top Chip

Next, we sought to incorporate vasculature into the system as we have with the other Organ-Chips we have developed with our proprietary technology^26^. As we have previously shown, the endothelium has important contributions in the development and functionality of the epithelial cells, including driving transcriptomic similarity to the tissue of origin^30^. To this purpose, the bottom channel of the Open-Top Chip, was seeded with tissue specific endothelial cells and then cultured under flow for 7 to 15 days as needed. At the end of the culture period the vascular channel was processed and stained for VWF, PECAM-1 (DC31) and VE-Cadherin. Perinuclear localization of VWF and homogeneous expression of the adhesion molecules, VE-Cadherin and PECAM-1 at the cell-periphery (Fig. 4; Fig. 5), indicate that endothelial cells form a tight monolayer and express markers of mature vasculature. These results corroborate the capability of the Open-Top Chip design to sustain the viability and differentiation of tissue specific endothelia in contact with its stromal element, similar to the tissue interactions *in vivo*.

**Figure 3.**
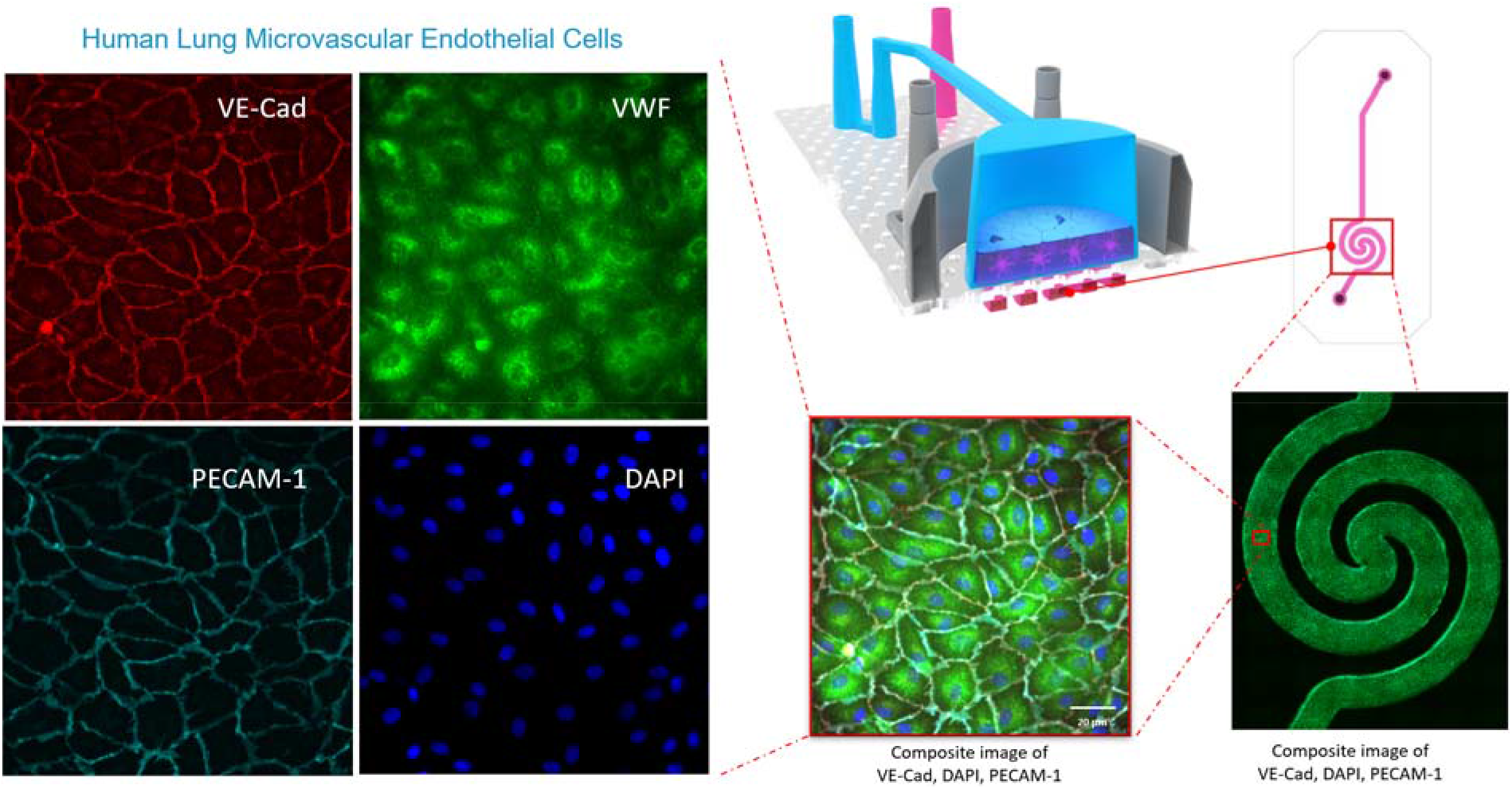
Open-Top Alveolus-Chip vascular compartment endothelial cell characterization: Immuno-fluorescence images showing endothelial marker expression and distribution along the vascular channel. It is lined with a tight monolayer of lung microvascular endothelial cells expressing endothelial specific markers PECAM-1 (cyan), VE-cadherin (red) and VWF (green).Composite image of the spiraled section of the bottom channel evidences that full cell coverage occurs homogeneously even in the section with complex geometry (spiral area).

**Figure 5.**
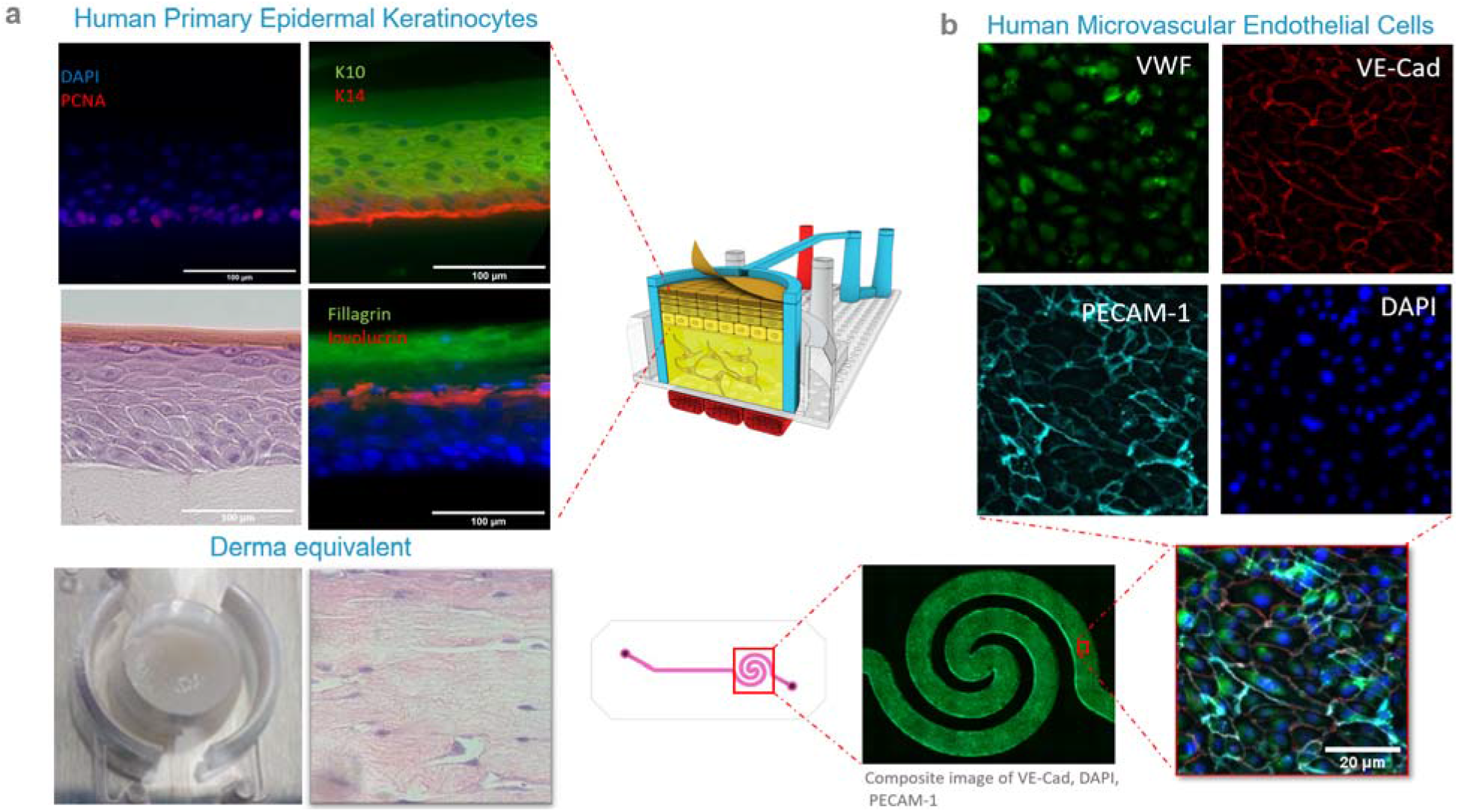
Open-Top Skin-Chip cell characterization: H&E cross section and immuno-fluorescence images showing: (a) epithelial marker of epidermis and fibroblast distribution across the tridimensional stromal compartment of the Open-Top Skin-Chip. (b) Endothelium coverage and endothelial markers of the vascular channel of the Open-Top Skin-Chip.

### Epithelial cell culture in the Open-Top Chip

Finally, we assessed how different epithelial cells respond and function in the Open-Top Chip when cultured in presence of stromal and vasculature components in the system, as described above. To this purpose we tested two different epithelial cell types: stratified keratinocytes, and alveolar epithelial cells. As described above, the epithelial channel has been designed to support establishment of ALI condition, that enables a more physiologically relevant mircroenvironement and has been shown to better support the differentiation and functional maturation of keratinocytes^31,32^, airway pneumocytes^33^ and alveolar cells^34,35,36^.

#### Skin epithelium model

In line with previous publications, we found that keratinocytes seeded on a fibroblast-embedded hydrogel coated with collagen IV (the main constituent of the basal lamina of the skin) and exposed to ALI for 15 days differentiate into a multi-stratified epithelium. Histology of the processed Open-Top Skin-Chip at this time point showed the presence of epidermal stem cells that were actively proliferating, as shown by PCNA positive cell nuclei. Expression of the basal cell marker keratin 14 was also found. In human tissue, the differentiation of keratinocytes from the basal to the spinous layer is characterized by the progressive shift in the expression of keratins, from keratin 14 to keratin 10. In addition, the positive expression of involucrin in the upper spinous layer and filaggrin in the cornified layer are indicative of a mature epidermal layer. In line with previous findings, histology of the Open-Top Skin-Chip showed the characteristic appearance of the stratified epidermis composed of cuboidal basal cells, squamous suprabasal keratinocytes and a superficial stratum corneum (Fig. 5).

#### Alveolus epithelium model

Similarly, alveolar pneumocytes grown on ECM composed of collagen IV, laminin and fibronectin expressed E-Cadherin at the cell-periphery, indicative of the formation of a tight monolayer. Alveolar type I cells stained positive for their specific markers for integral membrane protein HTI-56 and the mucin-type transmembrane T1-α/podoplanin. We also obtained positive staining for the lung apical plasma membrane protein, HTII-280, surfactant B and C, and key proteins involved in surfactant metabolism, including the lysosome-associated membrane glycoprotein 3, LAMP3, and the ATP binding cassette subfamily A member 3, ABCA3, all specific markers of alveolar type II cells. Importantly, the presence of alveolar surface microvilli was demonstrated by scanning electron microscopy (SEM). Histological analysis of sections from the Open-Top Alveolus-Chip confirmed the existence of two phenotypically distinct cell types: type I-like pneumocytes with the characteristic thin, squamous cell bodies and flattened nuclei, and type II-like pneumocytes with their large, cuboidal cell bodies (Fig. 6 a, b).

**Figure 6.**
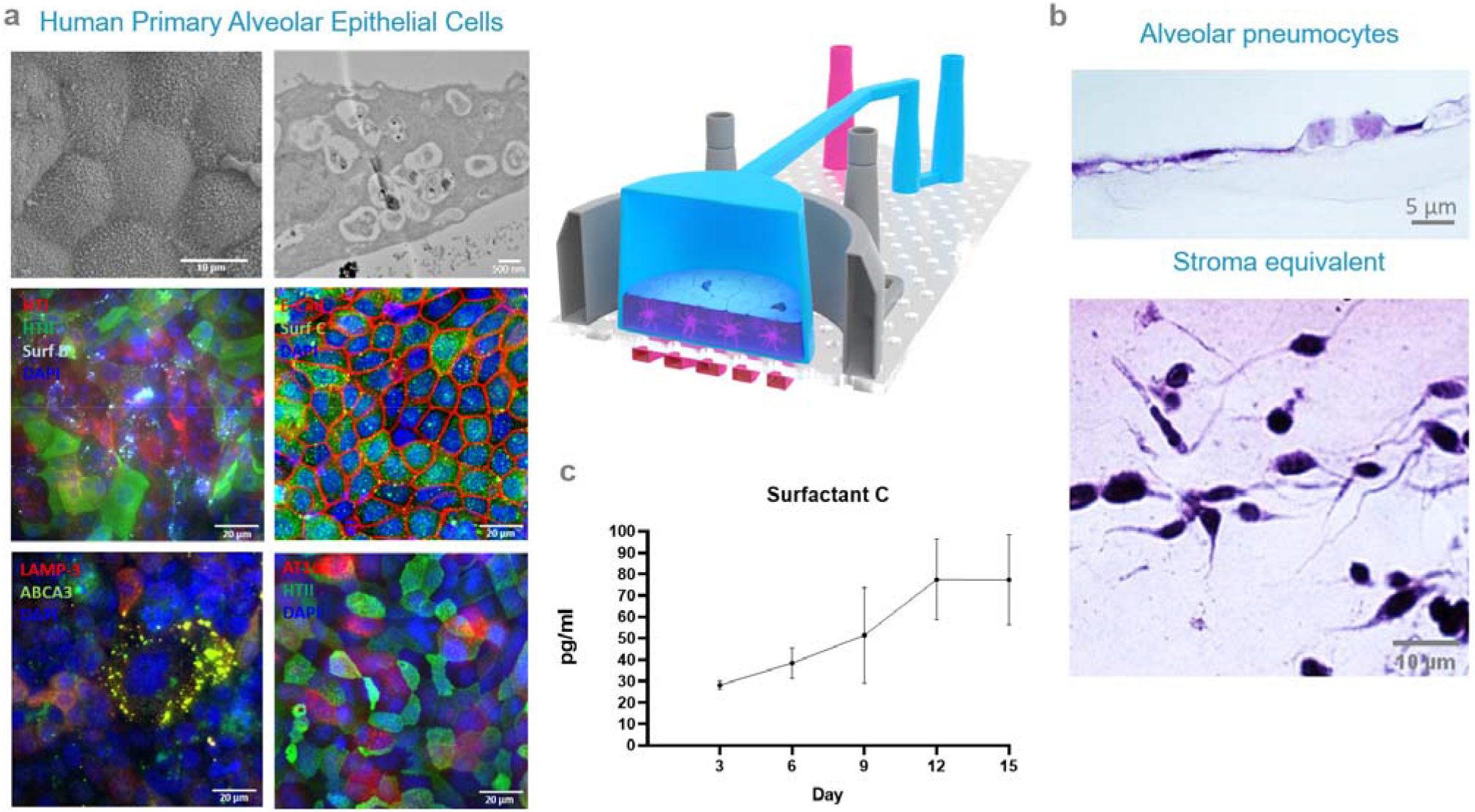
Open-Top Alveolus-Chip epithelial cell characterization: Scanning electron microscopy (SEM), transmission electron microscopy (TEM), immune-fluorescence and H&E cross section images showing: (a) epithelial marker presence and distribution of the alveolar epithelium across the surface of the epithelium compartment of the Open-Top Alveolus-Chip. (b) Histological images of the fibroblast distribution across the tridimensional stromal compartment and the alveolar epithelial cell morphology of the Open-Top Alveolar-Chip. (c) Surfactant C profile released by alveolar epithelium sampled at different time point during culturing in Open-Top Alveolus-Chip.

As described above, the Open-Top Alveolus-Chip was designed to recapitulate the native biomechanical environment of the human alveoli. The entire stromal compartment, including the inter-alveolar space and the epithelial monolayer, is surrounded by two vacuum channels that can be used to emulate the breathing motions, as previously published^37^. Upon application of mechanical stretching, we found enhanced expression of surfactant C in the stromal compartment, as has been reported with other systems^38,39^. Analysis of washes collected from the epithelial channel at different time points via ELISA, confirmed the secretion of surfactant C in soluble form (Fig. 6 c). TEM images also revealed the presence of lamellar bodies, associated with the production of surfactant C. Our findings show that human alveolar epithelial cells in the Open-Top Chip, grown on a stromal layer and exposed mechanical strain, expressed important tissue-specific makers of human alveolar cells and secreted surfactant C at measurable levels.

Next, we sought to evaluate whether this model enables stromal-epithelial interactions and captures the crosstalk between the three cell channels, as happens *in vivo*. We used an effective and well-studied stimulus, lipopolysaccharide (LPS), able to trigger an epithelial inflammatory response even in the absence of immune cells, as is the case of our model in its current version. Following treatment of the Open-Top Alveolus-Chip with LPS, we observed upregulation of the inflammatory surface marker ICAM-1 on endothelial cells, shown by immunohistochemistry, as well as increased levels of inflammatory mediators including IL-6, IL-8 and MPC-1, in the effluent of the vascular channel of the chip (Fig. 7). Notably, chips containing the vasculature alone, or vasculature and stroma but without epithelium, did not show an induction of ICAM-1 or increased cytokines release, in agreement with previous reports^40,41^. These data indicate functional interconnection between the three biological layers in the Open-Top Chip and confirms previous observations^4^. To further demonstrate the crosstalk between the alveolar epithelial layer and the vascular channel, we perfused the latter with whole blood containing CD41 labelled platelets for 15 min, while exposing the epithelium to LPS at a perfusion rate of 16.7μl/min (or 1ml/hr). We followed the movement of the labelled platelets and their interaction with the endothelial cells, by real time imaging. As it happens *in vivo*, inflammation in the vascular microenvironment, as simulated here by exposure to LPS, resulted in significant binding of platelets to the endothelial cells. The spatial distribution and localization of the platelets, as they adhere and aggregate at the endothelium is captured by live imaging (Movie 7.1). These findings show that the Open-Top Alveolus-Chip could be used to dissect cell-cell interactions in disease settings and assess drug toxicity and/or efficacy and even uncover donor-donor differences.

**Figure 7.**
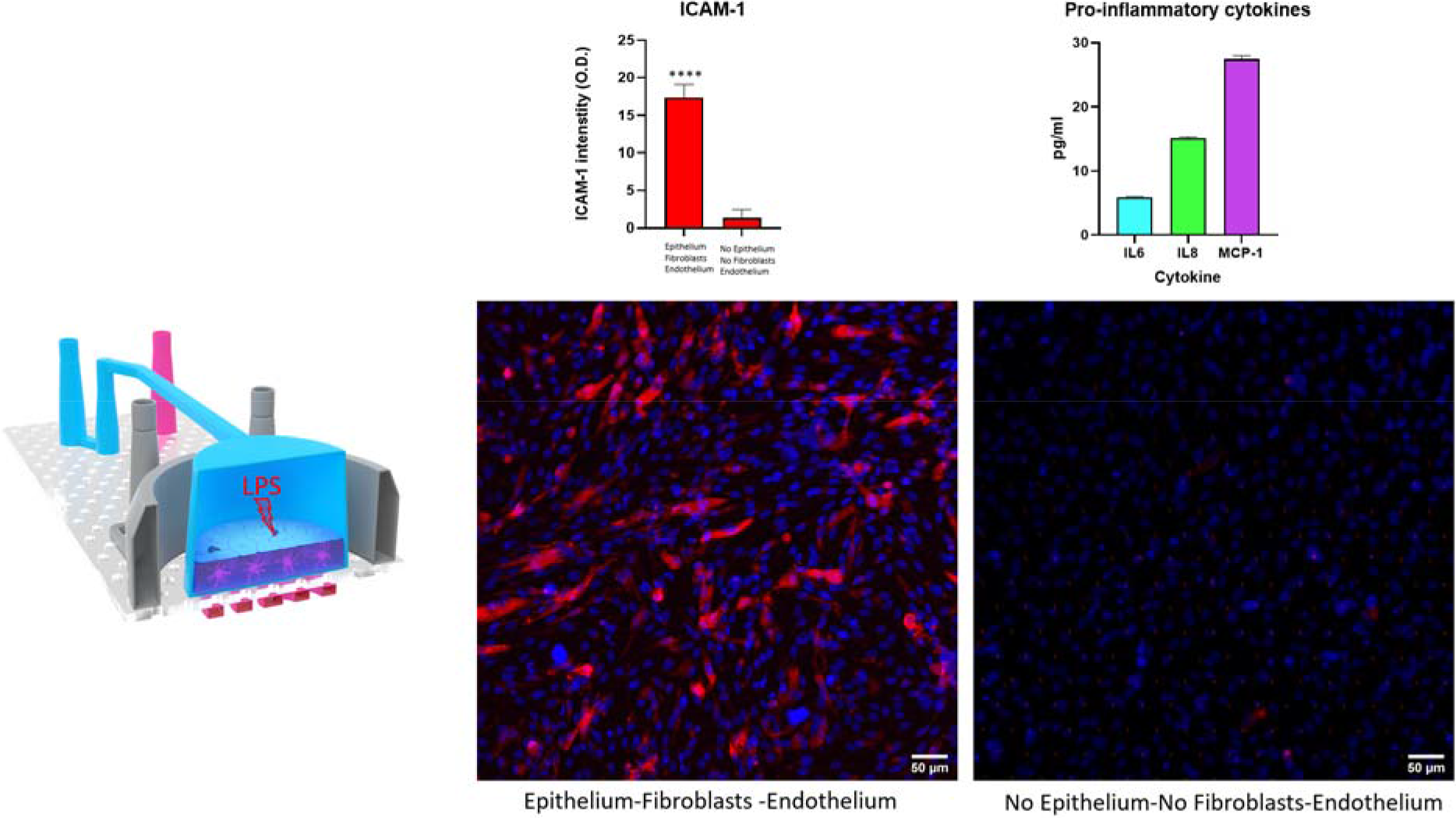
Graphs showing quantification of ICAM-1 intensity signal (left) and release of IL6, IL8 and MCP-1 cytokines (right) in response to LPS stimulation of the alveolar epithelial compartment. Immuno-fluorescence images showing ICAM-1 (red) expression in response to the stimulation with LPS. The induction of inflammation requires the presence of epithelium, fibroblasts, and endothelium to stimulate the maximum expression of ICAM-1 which is also sign of intimate interconnection between the three biological elements inside the chip. (Significance level p value: ****<0.0001)

## Discussion

Modeling the functions of the human epithelial tissues for the development of new therapeutics and furthering our understanding of disease mechanisms has been challenging, as these tissues are highly sensitive to the local microenvironment. Emerging evidence indicates the significance of tissue-tissue and cell-ECM interactions, as well as of the mechanical forces or shear stress the tissues are exposed to ^42,43^. Animal models have been instrumental in understanding the biological systems responses, but due to the intrinsic biological functional and genetic species differences, these finding often fail to translate to effective interventions for human patients^44^. On the other hand, traditional *in vitro* culture systems fail to reproduce the complex biological functions and interactions between the epithelia, parenchyma and vasculature required to faithfully model human biology and disease at the organ level^45^. The microengineered Organs-on-Chips technology is designed and engineered to address this translational gaps and enable more predictive models of the human biological responses *in vivo*^46^, as they provide fine control over the microenvironment and the ability to micro-fabricate complex structures at the tissue scale^47^.

Here we describe the design and engineering of the Open-Top Chip, a novel approach to recreate the epithelium-stroma-endothelial interactions, with the capability to control parameters of the microenvironment required to recapitulate *in vivo* relevant aspects of tissue functionality. In particular, leveraging microfabrication techniques, such as soft lithography and replica molding, we engineered a stretchable microfluidic chip made of a biocompatible, transparent, oxygen permeable, flexible elastomer (PDMS) to develop models of healthy and diseased human epithelia. The proprietary photoactivable heterobifunctional crosslinker used to functionalize the surface of the PDMS, resulted in a robust hydrogel-PDMS adhesion. The strength of the bond between the hydrogel and the PDMS is demonstrated through the ability to pneumatically stretch the Open-Top Chip without gel delamination, a common compromise of these systems. We designed a modular system, where tissue elements can be added or withdrawn to precisely characterize their role in health and disease states. For example, we were able to demonstrate the impact of the presence of epithelial cells and stromal fibroblasts on the differential expression of intracellular adhesion molecule-1 (ICAM1) in endothelial cells in response to LPS. The multilayered architecture of the chip design preserves the interaction between the epithelial and endothelial channels through diffusion of secreted molecules. To illustrate this, we induced inflammation in the Open-Top Alveolus-Chip by delivering LPS at the epithelial channel and then assessing its effects on the endothelial side. This experiment allowed us to define the contribution of the stroma in ICAM-1 expression in the endothelium. Specifically, LPS did not induce ICAM-1 expression in the absence of epithelium, even when fibroblasts have been incorporated in the stroma. These findings are of major significance for characterizing specific cell-to-cell interactions that can be targeted for therapeutic purposes and to enable better understanding of key processes and interactions driving disease mechanisms for new target identification and novel therapeutic development.

We next demonstrated that the endothelial channel can be perfused with whole blood under controlled conditions and using a well-established platelet adhesion assay we showed response of the endothelial cells to LPS, in line with *in vivo* findings. For example, we captured via high resolution videos, real-time platelet-vasculature interactions as shown by platelet-cell adhesion and aggregation. The dimensions of the Open-Top Chip are specifically designed for high-resolution imaging of living samples using conventional microscopy, and fluorescence, without compromising the sterility of the endothelium and epithelium. Here, we used high-resolution optical sectioning with confocal microscopy to visualize expression of specific endothelial cell surface markers. Quantitative characterization of the spatial and temporal distribution of immune cells recruited upon inflammatory activation is important in the study of infectious diseases. The Open-Top Chip is also compatible with recruitment assays of either PBMCs, or specific populations such as eosinophils or neutrophils, performed in microfluidics^48,49^ to model response to inflammatory stressors^49,50,51^.

The Open-Top Chip maintains the capability for independent treatments and sampling of the epithelial and endothelial channel through the top and bottom microchannels, similar to Emulate’s other chip designs. Medium effluent or other secreted fluids collected by washing from epithelial channel (mucus, surfactant) and/or vascular washing can be sampled over time to assess secretory activity, directional transport, uptake and dynamic metabolism of compounds for pharmacodynamic studies in more physiologically relevant conditions than currently supported by the existing systems. The potential for real-time, non-invasive visualization of the biological features of the recreated organ unit without sacrificing its sterile environment, together with evaluation of secreted factors, enabled us to obtain proof-of-concept quantitative data illustrating *in vivo* relevant responses of the cells. Future studies could expand the disease-relevant repertoire of available endpoints to define individual-specific efficacy and toxicity, via parallel clinical trials on-Chip.

In summary, we describe the development of a microengineered system for better simulating the human epithelium-stroma-endothelial interactions and their potential applications in modeling complex human biology and disease. The Open-Top Chip may be leveraged for functional assessment of pharmacological compounds targeting the stroma, a tissue of increasingly recognized contribution in the progress of a number of diseases such as inflammatory and infectious diseases, tissue fibrosis and cancer.

## Supporting information

Movies (stretching)

---

**Figure S1.**
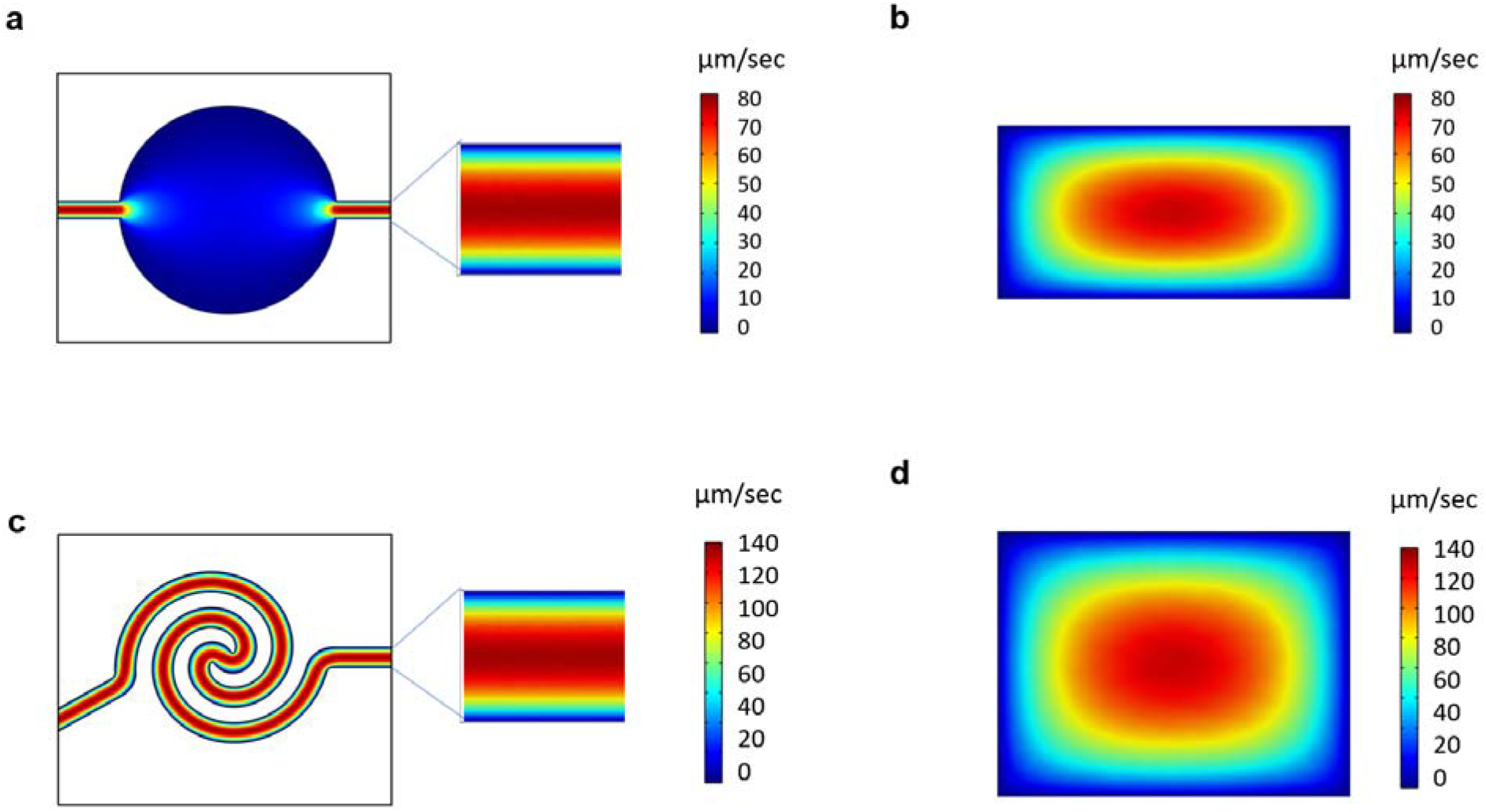
COMSOL Multiphysics modeling of top and bottom channel fluid flow profiles: (a) Top view of well-developed, laminar flow velocity profile in top channel (200μm(height)×600μm(width)). (b) Cross-sectional view of top channel velocity profile. (c) Top view of well-developed, laminar flow velocity profile in bottom spiraled channel (400μm (height) ×600μm(width)). d) Cross-sectional view of bottom channel velocity profile.

**Movie S2.1.**
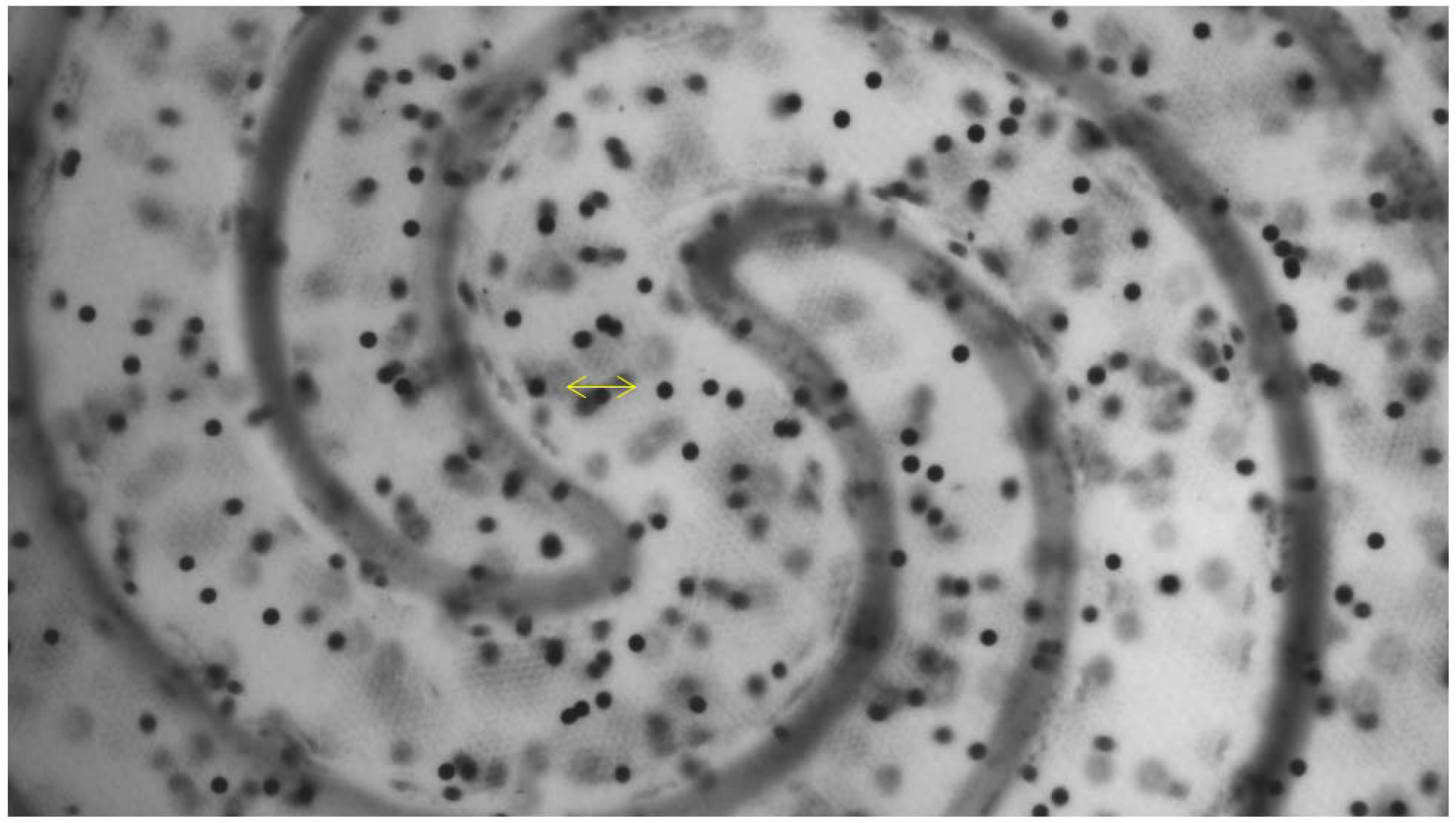
Video showing the displacement of bead inside the gel when negative pressure is applied to the two curved vacuum-channels. By measuring the displacement between beads, we estimate the percentage of strain that cells experience on-chip when breathing motion is applied.

**Movie S2.2.**
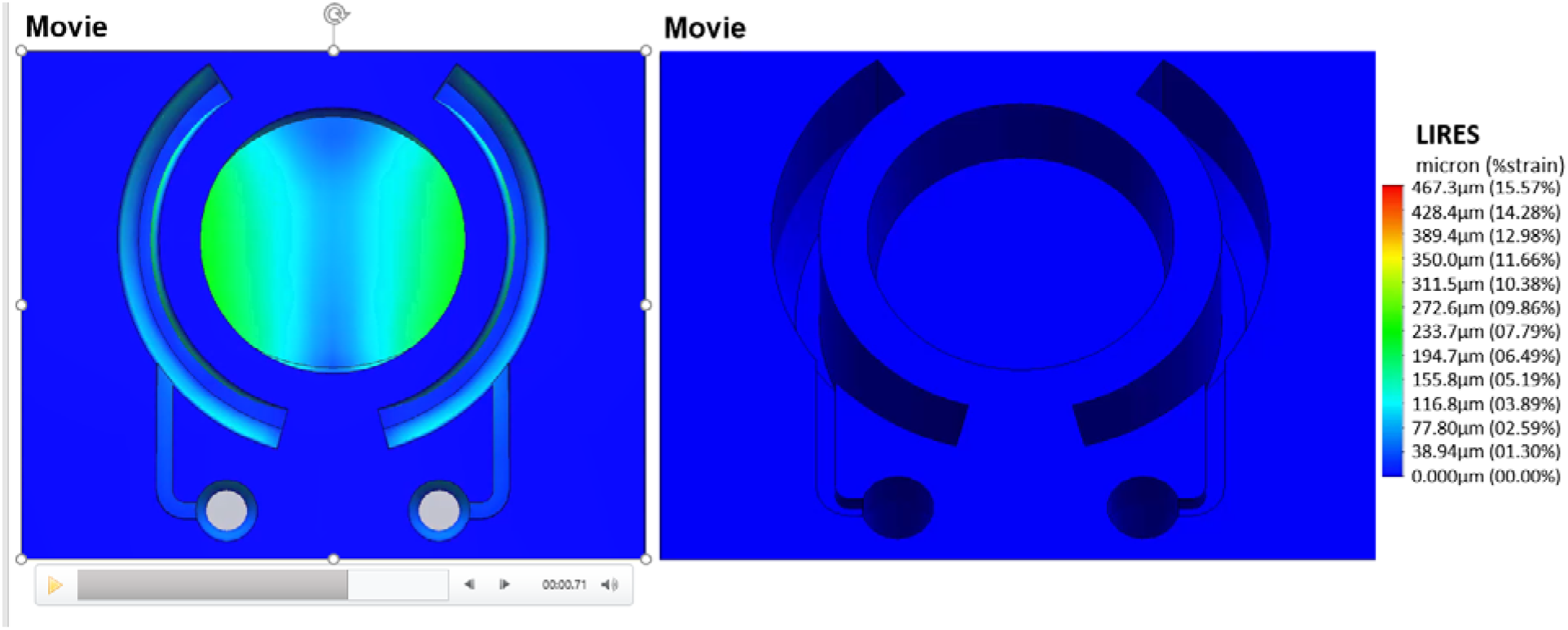
Video showing the Finite Element Analysis (FEA) model of the applied mechanical stretching along the surface of the stroma equivalent (top and angled view).

**Movie 7.1.**
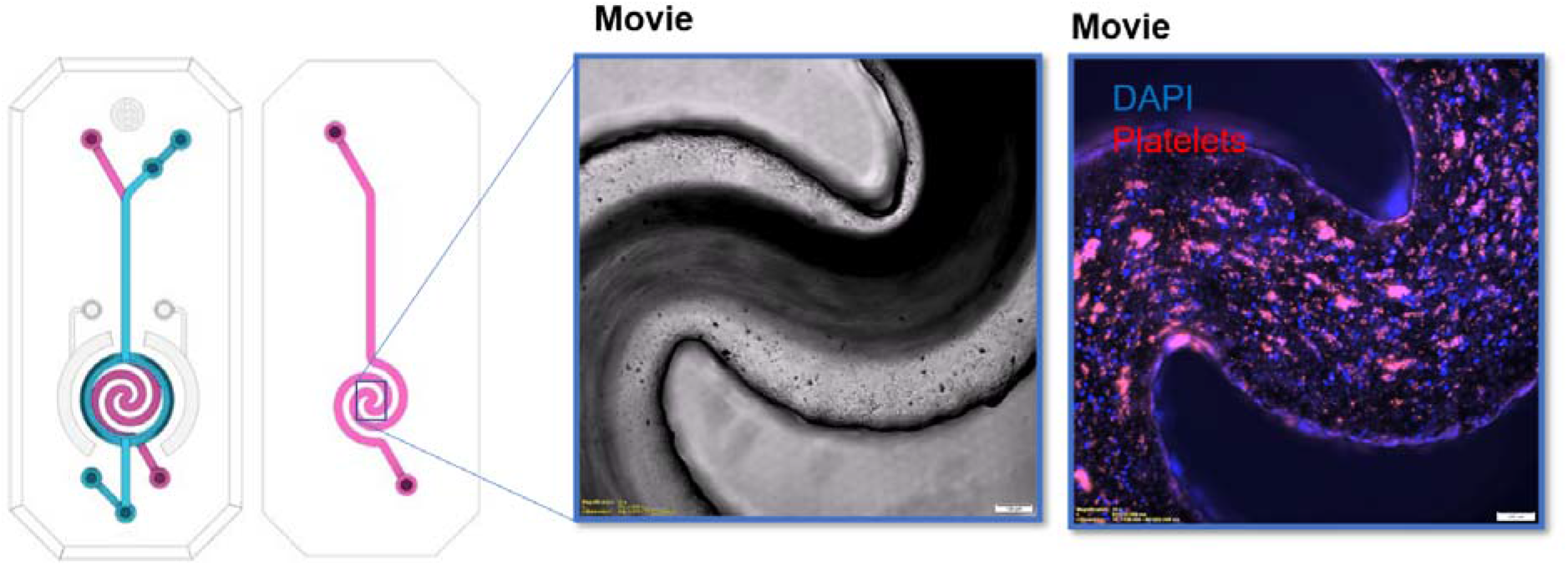
Videos in phase contrast (right) and fluorescence (left) showing blood perfusion and temporal and spatial platelet adhesion to the inflamed endothelium in response to LPS stimulation of the alveolar epithelial compartment.

