## Supplementary material for "“A novel Organ-Chip system emulates three-dimensional architecture of the human epithelia and allows fine control of mechanical forces acting on it.”": Movies (stretching)

### Slide 1
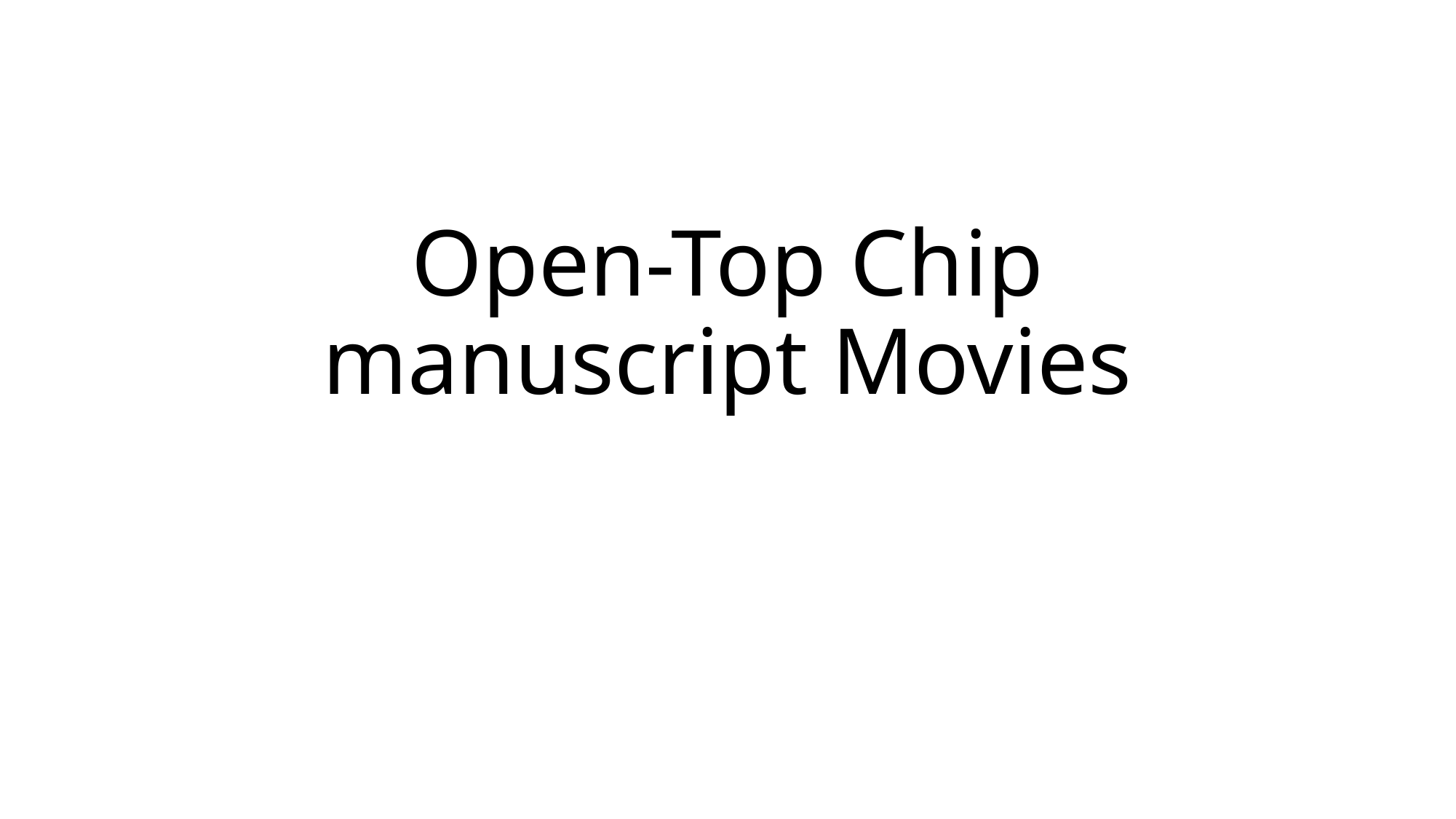

#### Open-Top Chip manuscript Movies

### Slide 2
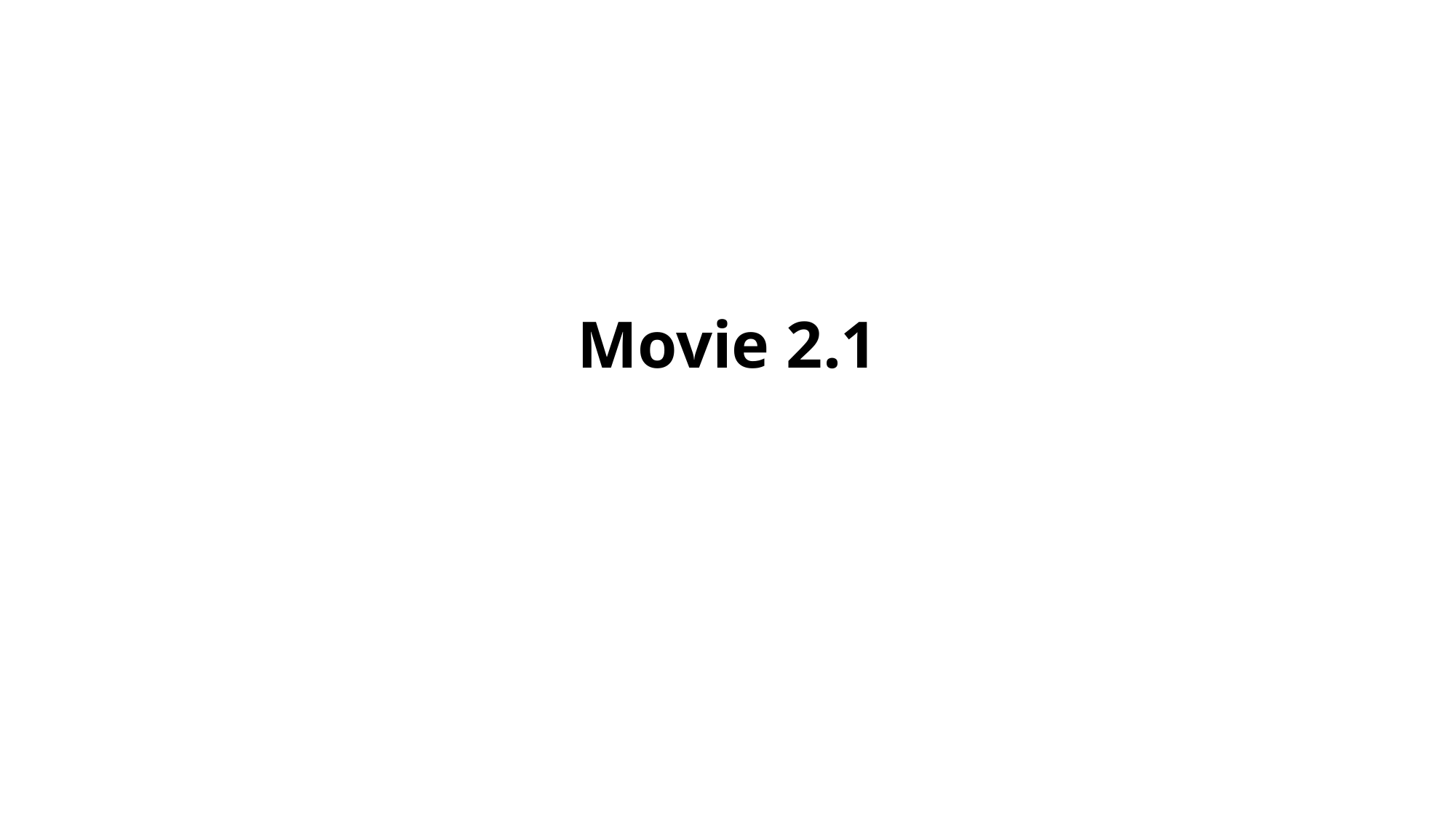

#### Movie 2.1

### Slide 3
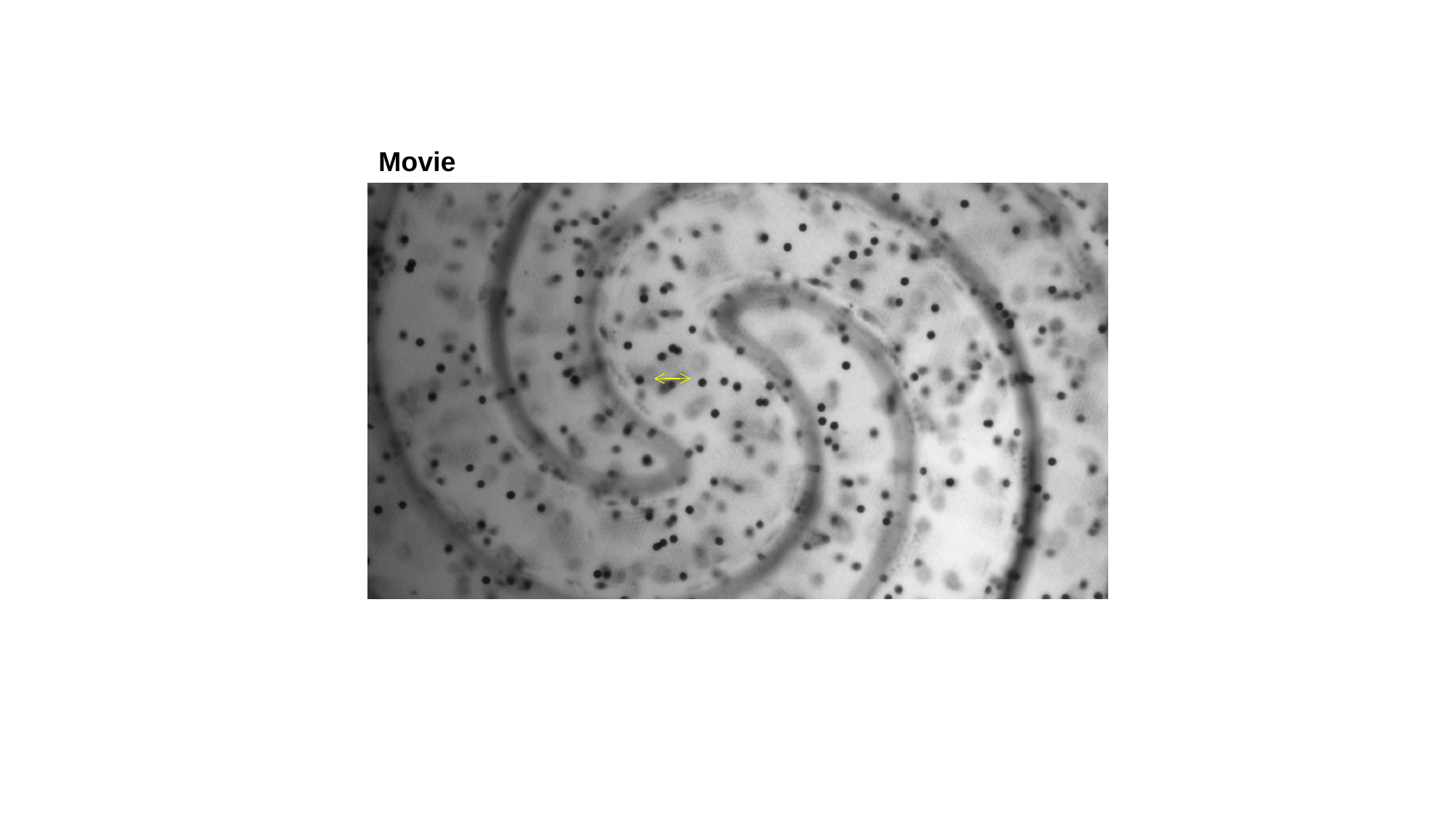

Movie

### Slide 4
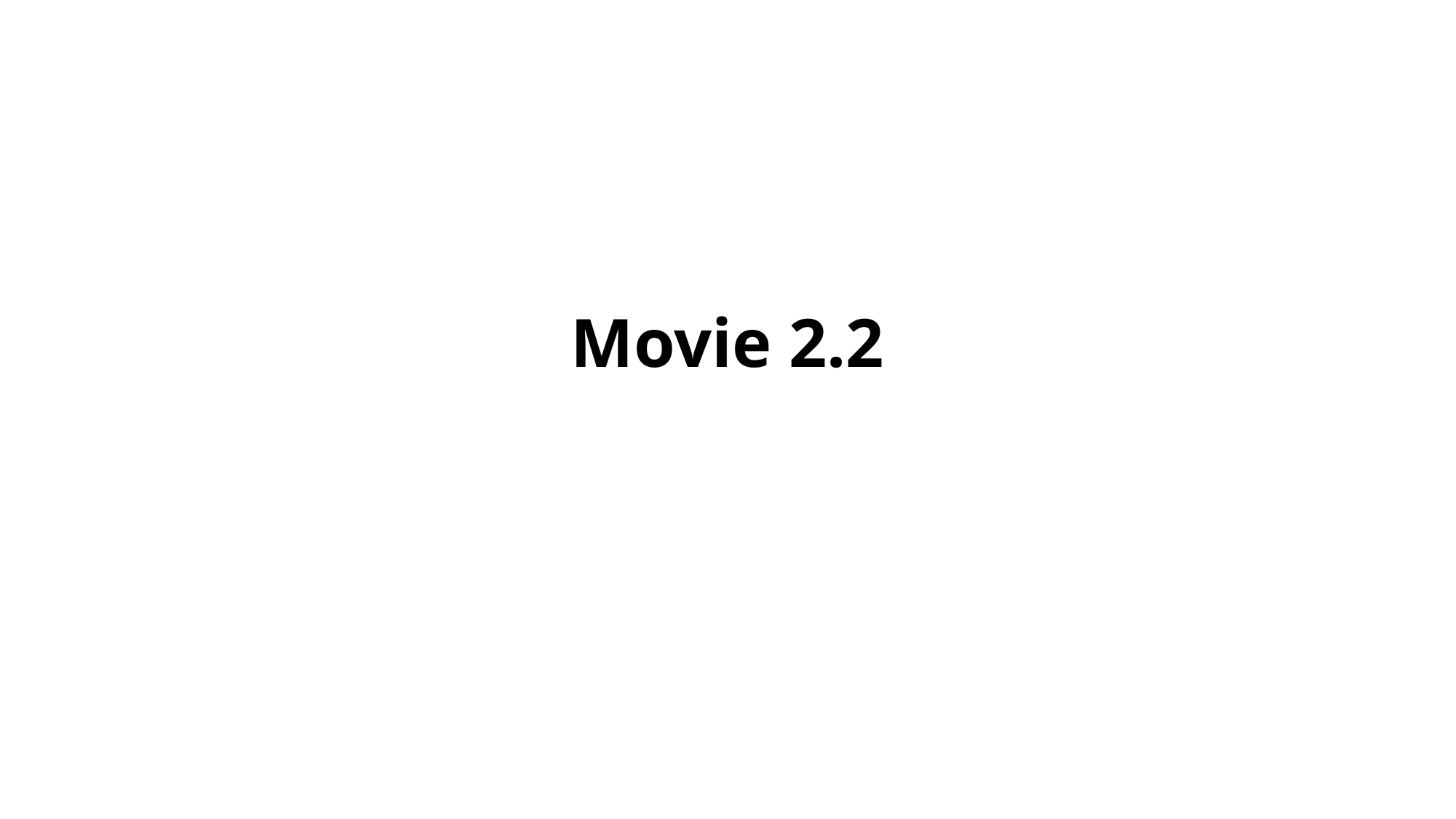

#### Movie 2.2

### Slide 5
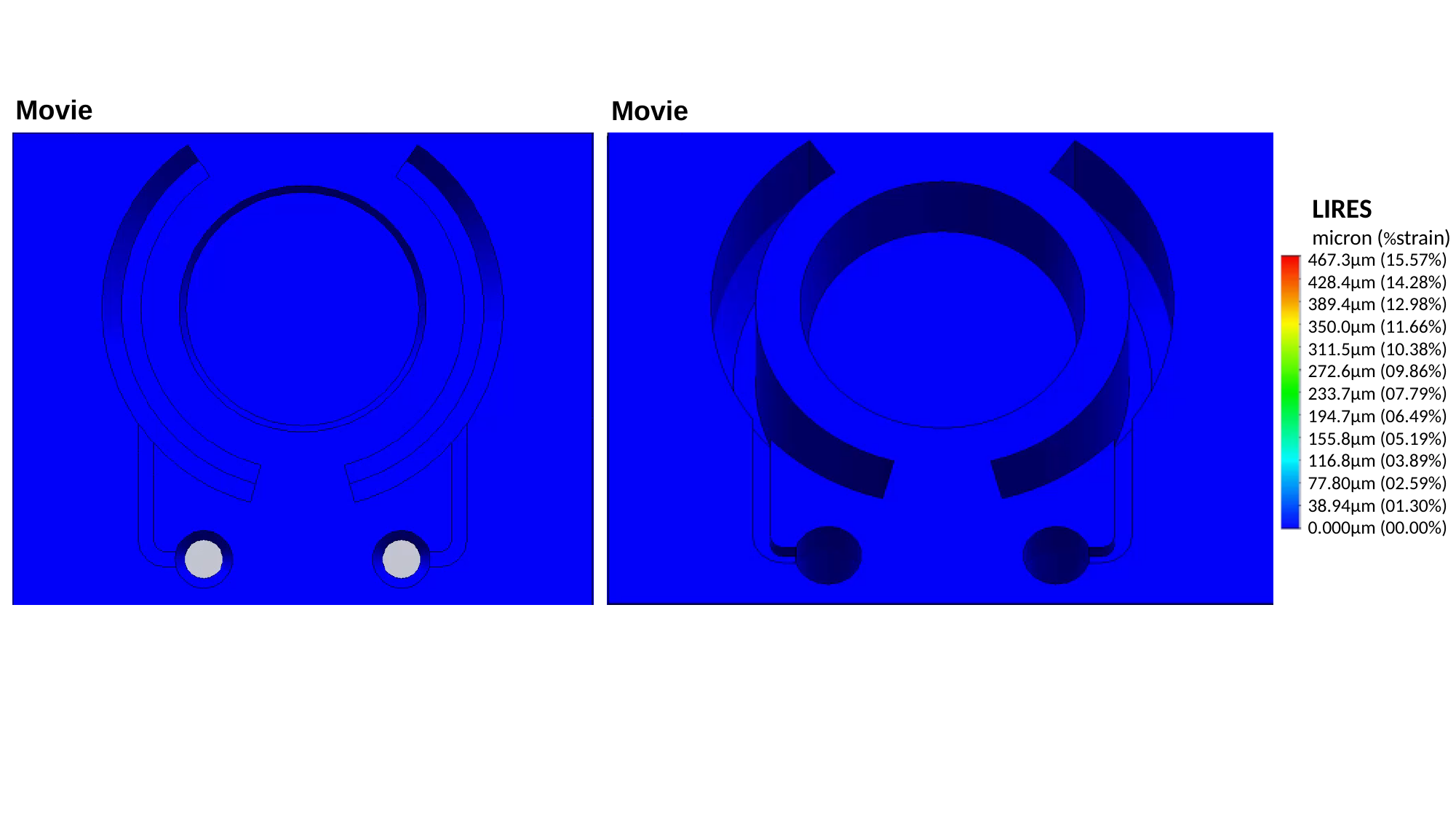

Movie
Movie
LIRES
micron (%strain)
467.3µm (15.57%)
428.4µm (14.28%)
389.4µm (12.98%)
350.0µm (11.66%)
311.5µm (10.38%)
272.6µm (09.86%)
233.7µm (07.79%)
194.7µm (06.49%)
155.8µm (05.19%)
116.8µm (03.89%)
77.80µm (02.59%)
38.94µm (01.30%)
0.000µm (00.00%)

### Slide 6
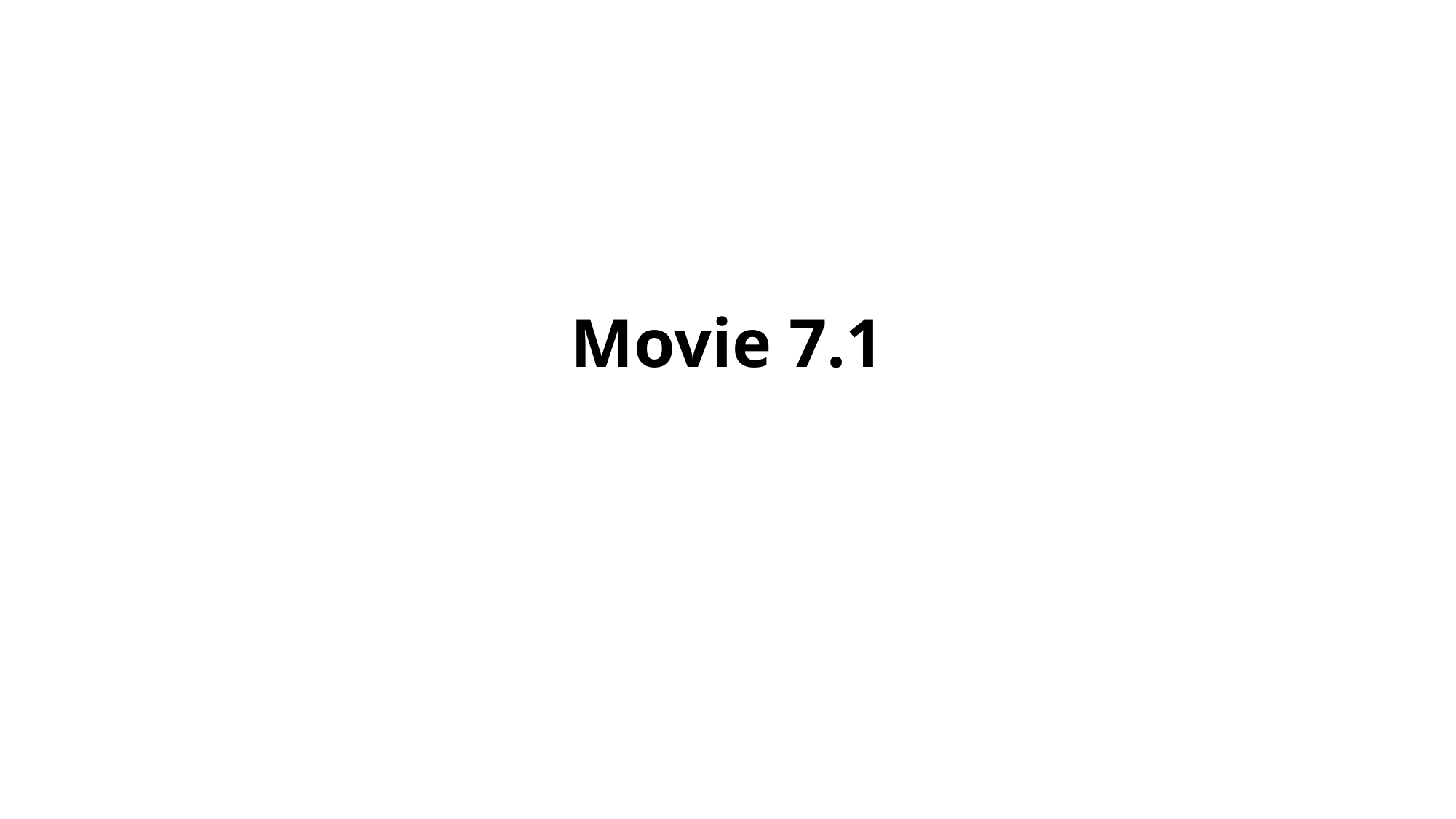

#### Movie 7.1

### Slide 7
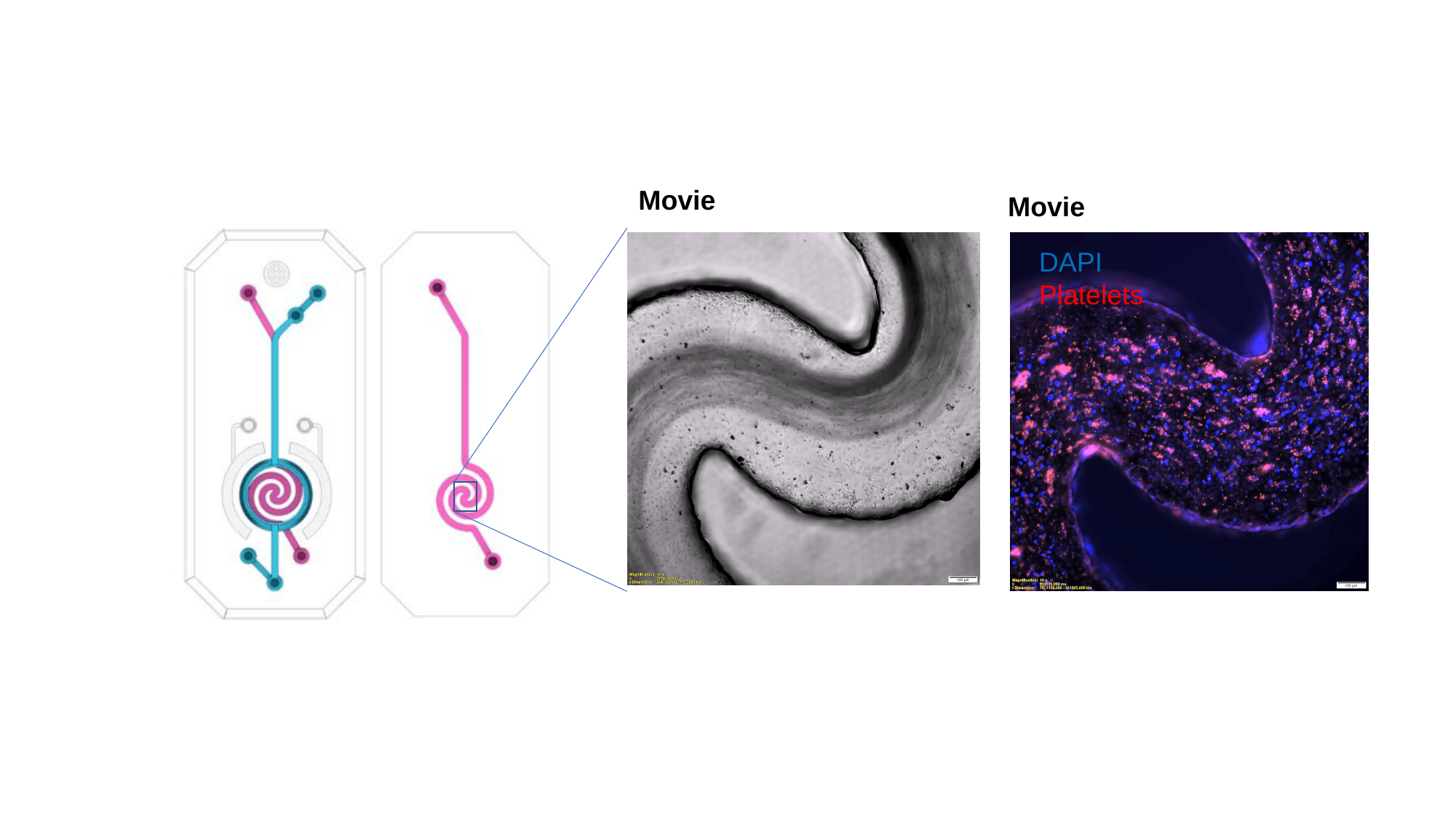

Movie
Movie
DAPI
Platelets
